# Single-molecule imaging reveals the coupling of translation and mRNA decay

**DOI:** 10.1101/2021.06.07.447377

**Authors:** Pratik Dave, Esther Griesbach, Gregory Roth, Daniel Mateju, Jeffrey A. Chao

**Affiliations:** Friedrich Miescher Institute for Biomedical Research, 4058 Basel, Switzerland

## Abstract

The relationship between mRNA translation and decay is incompletely understood, with conflicting reports suggesting that translation can either promote decay or stabilize mRNAs. The effect of translation on mRNA decay has mainly been studied using ensemble measurements and global inhibitors of transcription and translation, which can mask the underlying mechanisms. We developed a single-molecule imaging approach to control the translation of a specific transcript that enabled simultaneous measurement of translation and mRNA decay. Our results demonstrate that mRNAs undergoing translation are degraded faster than non-translating ones, although with slower kinetics than translation-coupled degradation of transcripts targeted by NMD. Furthermore, our results indicate that miRNAs mediate efficient degradation of both translating and non-translating target mRNAs. Single-molecule measurements of translation and decay reveal a predominant role of mRNA decay in miRNA-mediated regulation. Simultaneous visualization of translation and decay on single mRNAs provides a framework to study how these processes are interconnected in cells.

## INTRODUCTION

When mRNAs enter the cytoplasm, they are translated into proteins and eventually targeted for mRNA decay. The balance between these two processes is critical to establish cellular homeostasis. While the steady state levels of mRNAs is determined by the balance between the transcription and decay rates (Dolken et al., 2008; Elkon et al., 2010), the dynamic changes in RNA levels as a response to external stimuli were found to be dependent on RNA turnover alone (Kawata et al., 2020). There are several intrinsic cis-acting elements within the mRNA sequence that can influence its stability. These factors include codon-optimality (Buschauer et al., 2020; Presnyak et al., 2015; Wu et al., 2019), mRNA modifications (Herzog et al., 2017), AU-rich elements or miRNA binding sites in the 3’UTR (Garneau et al., 2007), GC-composition (Courel et al., 2019) and the sequence of 5’UTR (Jia et al., 2020).

The stability of mRNAs is also influenced by *trans*-acting factors that associate with an mRNA and its interaction with the ribosome is thought to play central role. While the connection between translation and mRNA decay has long been appreciated, a consensus on whether ribosomes increase or decrease mRNA stability has not yet been formed (Heck and Wilusz, 2018). An early experiment linking translation to mRNA decay came from studying c-myc mRNA degradation, almost three decades ago, where they found that shortening of the poly(A)-tail was translation-dependent (Laird-Offringa et al., 1990). More recently, genome-wide studies have shown that mRNA decay can occur simultaneously on actively translating mRNAs in yeast and mammals (Pelechano et al., 2015; Tuck et al., 2020). This co-translational decay model is further supported by a growing number of decay factors that directly associate with ribosomes (Buschauer et al., 2020; D’Orazio et al., 2019; Ibrahim et al., 2018; Juszkiewicz et al., 2018; Singh et al., 2020; Tesina et al., 2019; Tuck et al., 2020). It is, however, not clear if co-translational decay of mRNAs is a direct consequence of the process of translation itself. In fact, other studies have indicated that translation protects the mRNAs from being degraded (Chan et al., 2018; Jia et al., 2020), and that mRNA translation and decay are mutually exclusive processes (Brengues et al., 2005).

Previously, we found that inhibiting translation at initiation (eIF2α phosphorylation), elongation (cycloheximide and harringtonine) or causing premature termination (puromycin) resulted in mRNA stabilization. While these results suggested that translation promoted mRNA decay, we could not exclude the possibility that global translation inhibition confounded the effect by altering other pathways (Gowrishankar et al., 2006; Hilgers et al., 2006; Muckenthaler et al., 1997; Santos et al., 2019; Yamagishi et al., 2014). Importantly, ensemble measurements that have also studied the interplay between translation and mRNA decay are inherently limited by normalization of the data between different types of experiments (e.g. ribosome profiling and RNA-seq or luciferase assays and RT-PCR) and the sensitivity of these techniques.

Here, we describe a single-molecule methodology that combines transcript-specific regulation of translation by the iron response element and TREAT to study the coupling of translation and decay in single cells. Using this system, we observe enhanced degradation of transcripts undergoing translation, directly demonstrating that association with the ribosome promotes mRNA decay. This translation-dependent decay, however, occurs with slower kinetics than degradation of transcripts targeted by the NMD pathway indicating this approach enables different translation-dependent mRNA decay mechanisms to be characterized. Interestingly, when a miRNA-binding site was introduced in the 3’UTR of a transcript, these mRNAs were rapidly degraded regardless of their translation status. Furthermore, by combining SunTag and TREAT imaging, we find that miRNAs primarily function by accelerating mRNA decay and that inhibition of translation has little, if any, contribution to repression. Our study provides insights into the connection between translation and mRNA decay and a framework for further characterizing their interplay in single cells.

## RESULTS

### Single-molecule imaging of iron-inducible translation

To characterize the relationship between translation and mRNA decay, a reporter RNA was designed that allows us to control the translation of a specific transcript without the pleiotropic effects of inhibiting global mRNA translation. The reporter mRNA contains the 5’UTR from ferritin heavy chain mRNA, which in turn contains an Iron Responsive Element (IRE). Under low cellular iron concentrations, IREs recruit Iron regulatory proteins (IRP1/2) that prevents translation initiation by blocking the recruitment of pre-initiation complex (Hentze et al., 1987a; Hentze et al., 1987b; Muckenthaler et al., 1998) (Figure 1A). With increasing iron levels, the IRP1/2 proteins dissociate from the IRE enabling translation initiation. In order to achieve functional repression, it is critical that IRE stem-loop is positioned no further than 60 nucleotides from the cap, beyond which mRNAs lose translation repression and become constitutively active (Goossen and Hentze, 1992). For this reason, the IRE was placed 31 nucleotides from the 5’cap in our reporter.

**Figure 1.**
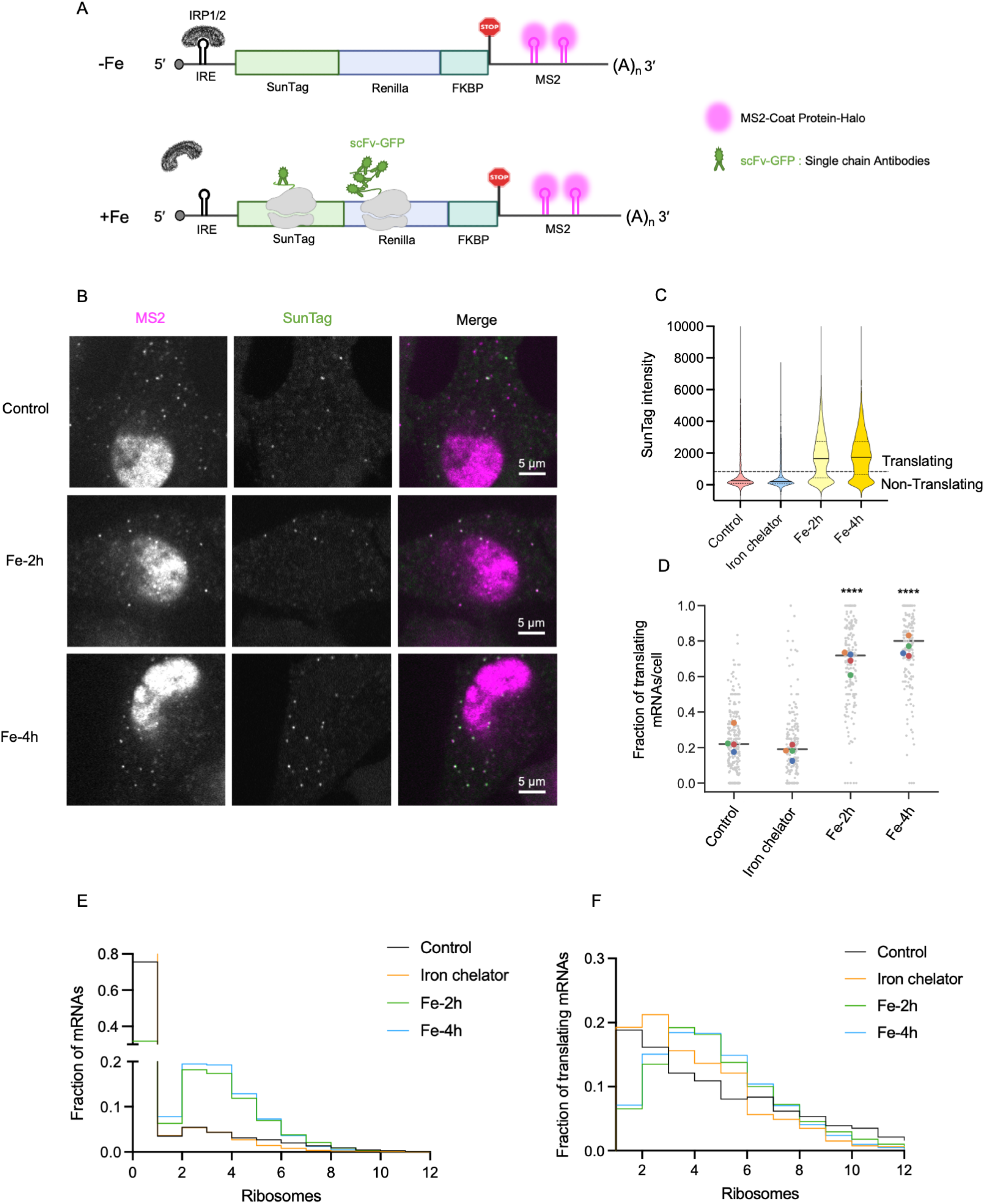
Single-molecule imaging of Iron-inducible mRNA translation. **A.** Design of the translation-inducible IRE-SunTag reporter used in the current study. The 5’UTR of human Ferritin heavy-chain mRNA containing the Iron Responsive Element (IRE) recruits IRP1/2 proteins in absence of Fe. Upon high Cellular Fe concentration, the IRP1/2 dissociate from IRE, facilitating ribosome recruitment. The reporter was stably integrated in HeLa-11ht cells expressing scFV-GFP and MCP-Halo. **B.** Live-cell single-molecule imaging of IRE-SunTag reporters in absence iron and after treatment with 150 μM ferrous ammonium citrate as iron source for 2h (Fe-2h) or 4h (Fe-4h). Magenta spots represent mRNAs and green spots represents SunTag indicating translation. **C.** Quantification of the SunTag intensities corresponding to the IRE-SunTag reporter mRNAs in absence of Fe (Control-5420 mRNAs), in presence of Iron chelator (4620 mRNAs), in presence of Fe treated for 2h (Fe-2h-3140 mRNAs) or Fe treated for 4h (Fe-4h-3688 mRNAs), from 4 independent experiments. The background GFP intensity of the cell was subtracted from the measured SunTag intensity at each mRNA. The dotted line represents the cut-off based on intensity of single SunTag polypeptide. **D.** The distribution of fraction of translating mRNAs per cell in Control (253 cells), Iron chelator (200 cells), Fe-2h (196 cells) and Fe-4h (197 cells) from 4 independent experiments. **** indicates p-value < 0.0001 when compared to Control. Color-coded dots represent averages from individual experiment set. **E and F.** Quantification of number of ribosomes on mRNAs represented in figure 1C. Y-axis represents relative fraction of mRNAs from 4 independent experiments. The frequency of mRNAs with number of ribosomes for all the mRNAs (E) or only the translating mRNAs (F) is represented.

To visualize the translation of single RNA molecules in living cells, the SunTag cassette was integrated into the coding sequence as an N-terminal fusion protein with Renilla luciferase (Pichon et al., 2016; Wang et al., 2016; Wu et al., 2016; Yan et al., 2016) and MS2 stem-loops were incorporated into the 3’UTR. The SunTag cassette encodes for an array of antigenic epitopes allowing visualization of translation of the reporter mRNA via detection of nascent polypeptide in the cells expressing the cognate single-chain antibodies fused to GFP (scFV-GFP). The 24 MS2 stem-loops are bound by an MS2 coat protein (MCP)-Halo fusion protein that enables visualization of single mRNAs. This reporter was integrated under a doxycycline-inducible promoter into a single genomic locus in a previously established HeLa cell line stably expressing scFV-GFP antibody and MCP-Halo (Mateju et al., 2020; Voigt et al., 2017; Wilbertz et al., 2019). In order to limit the accumulation of mature SunTag peptides in the cytoplasm that can increase the background signal, the destabilized FKBP domain was fused in the C-terminus of the SunTag-Renilla (Banaszynski et al., 2006).

To test the functionality of the reporter, live-cell imaging of the IRE-SunTag reporter was performed in either the absence or presence of iron. In untreated cells, the majority of transcripts were not actively translating as single mRNAs did not have a detectable SunTag signal (Figure 1B, Video S1). Since ferric ammonium citrate is converted intracellularly to Fe^2+^ and prolonged exposure results in its export, a time course was performed to determine an appropriate imaging window. After treatment of cells with ferric ammonium citrate (150 μM) for either two or four hours, most mRNAs were now found to be translating (Figure 1B, supplementary Video S2 and S3), while a one-hour time point resulted in few translating mRNAs (data not shown). It should be noted that a small fraction of mRNAs was determined to be translating even in untreated cells (figure 1B, supplementary Video S1) as well as in cells treated with an iron chelator (supplementary Video S4), which indicated that the IRE does not completely inhibit translation of all mRNAs.

In order to quantify the fraction of mRNAs translating in untreated and iron-treated cells, a high-throughput image-analysis workflow was developed. mRNAs were tracked using single-particle tracking of MCP-Halo spots and the corresponding intensities of the SunTag were quantified at the same coordinates. The background GFP-intensity was subtracted from the mean SunTag intensity of the individual tracks. To assign the translation status of mRNAs, the intensity of single mature SunTag peptides in puromycin-treated cells was quantified and used as a cutoff to determine translation status of an mRNA as (Figure S1). This approach allowed us to quantify the translation status of thousands of IRE-Suntag reporter mRNAs in living cells that were untreated, iron-treated and iron-chelator treated and revealed the heterogeneity in translation of individual mRNA molecules (Figure 1C). The average fraction of translating mRNAs was ~20% mRNAs in untreated and iron chelator-treated cells and this increased to ~70-75% in iron-treated cells (Figure 1D). As a control, translation of a reporter lacking the IRE in the 5’UTR (ΔIRE-SunTag) was also quantified (Figure S2A and S2B, supplementary Video S5 and S6). For the ΔIRE-SunTag reporter, the distribution of SunTag intensities as well as the fraction of translating mRNAs per cell remained unchanged upon treatment of cells with iron as compared to the untreated control, demonstrating the specific activation of translation of IRE-SunTag mRNAs (Figure S2C and S2D).

While the IRE is well-known to inhibit translation, the distribution of SunTag intensities enables its regulatory mechanism to be further characterized in cells. Based on the intensity of a single SunTag peptide, the ribosome occupancy on individual RNAs was quantified. If all mRNAs are analyzed, we find that most mRNA are not actively translating in untreated or iron-chelator treated cells and a small fraction are in polysomes containing 2-8 ribosomes (Figure 1E, S3A). Interestingly, if we only analyze the ribosome occupancy of translating mRNAs in untreated, iron-treated and iron-chelator treated cells, we observe that all conditions contain a similar distribution (Figure 1F) and the average number of ribosomes per mRNA remained similar (Figure S3B). This data thus suggests that mRNAs undergoing translation in untreated cells have similar initiation rates as translating mRNAs in iron-treated cells and that the basal expression level observed for IRE-mediated regulation in untreated cells comes from a few mRNAs that have escaped IRP1/2-IRE mediated repression (“on”-state mRNAs), whereas the majority of mRNAs remain bound by IRP1/2 proteins and are repressed (“off”-state mRNAs).

### mRNA decay kinetics of translating and non-translating mRNAs in single cells

Having established an inducible system to control the mRNA translation of a specific transcript, we next applied this tool to study the kinetics of mRNA decay at the single-molecule level in the presence and absence of translation. To achieve this, the IRE was incorporated in the 5’UTR of a previously established TREAT (3(*t*hree)’-*R*NA *e*nd *a*ccumulation during *t*urnover) reporter, which allows us to distinguish between an intact mRNA and a stabilized 3’degradation intermediates using single-molecule fluorescence in situ hybridization (smFISH) due to the presence of tandem XRN1-resistant pseudo-knots (PKs) (Figure 2A) (Horvathova et al., 2017). The IRE-TREAT reporter RNA was integrated into a single genomic locus in HeLa cells under the control of a doxycycline-inducible promoter. The resulting IRE-TREAT expressing cell line allowed us to study the effect of translation on the degradation of mRNAs in single cells.

**Figure 2.**
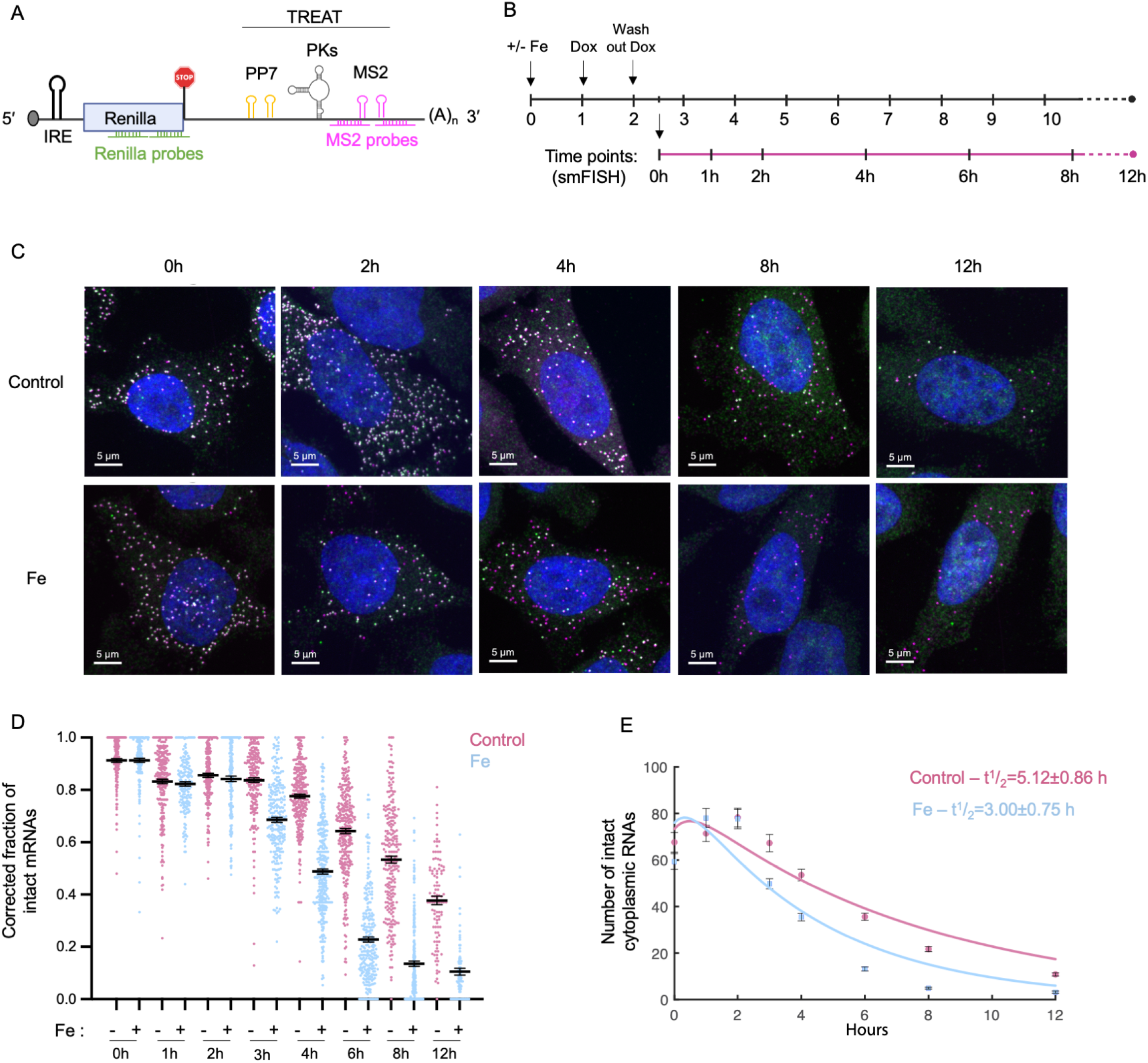
IRE-TREAT reporter mRNA to study the decay kinetics in presence or absence of translation. **A.** Schematic representation of the IRE-TREAT reporters used to study mRNA decay kinetics in presence and absence of translation. The 5’UTR contains the IRE and the 3’UTR contains the TREAT cassette (as described in (ref)) consisting 24x PP7 stem-loops, 2x-Xrn-1 resistant pseudoknots (PKs) and 24x-MS2 stem-loops in that order. The 5’UTR contain the IRE, allowing inducible translational control. smFISH probes targeting Renilla in ORF (upstream PKs) and MS2 stem-loops in 3’UTR (downstream PKs) is indicated. The reporters were integrated in HeLa cells under doxycycline inducible promoter. **B.** Schematic representation of the experiment design to study the mRNA decay kinetics in presence or absence of translation. The cells were pre-treated with 150 μM ferrous ammonium citrate for 1h (untreated for control samples) before induction of transcription by doxycycline for 1h. Cells were thoroughly washed to remove doxycycline and stop the transcription and were subsequently maintained in DMEM with or without iron. 30 minutes after the wash, cells were fixed and processed for smFISH at indicated time intervals. **C.** Representative smFISH images of cells expressing IRE-TREAT reporters in presence or absence of iron at indicated time-points. Magenta spots represent MS2 and green spots represent Renilla. Dual labelled spots indicate the intact reporter mRNAs and single labelled MS2 spots represent stabilized intermediates indicating degraded mRNA. **D.** Quantification of mRNA decay from the smFISH images. Y-axis represents fraction of intact mRNAs corrected for the detection efficiency for the smFISH probes (see supplementary figure S4). Each dot represents fraction of intact mRNAs per cell for 2 independent experiments. Mean and error bars indicating SEM is represented. Peach dots represent control and blue dots represent iron-treated cells. **E.** Quantification of number of cytoplasmic intact RNAs in control and Fe-treated cells. Mean and error bars representing SEM is indicated. A mathematical model was fitted to the data (solid lines) to obtain the degradation rates. The half-life with standard deviation is indicated.

To study the decay kinetics of the IRE-TREAT reporter, a pulse (1h) of doxycycline was used to induce transcription of the reporter and cells were fixed at the indicated time-points and processed for smFISH (Figure 2B). Cells were pre-treated with iron for 1h prior to induction of transcription for iron-treated samples. To distinguish between intact and degraded RNA fragments in fixed cells, smFISH was performed using fluorescently-labelled probes targeting Renilla luciferase in the coding sequence (upstream of PKs) and the MS2 stem-loops in the 3’UTR (downstream of PKs) (Figure 2A).

At the initial 0h time-point, the majority of the IRE-TREAT mRNAs in both untreated and iron-treated cells were found to be intact, as observed by dual labelled RNA spots (Figure 2C). mRNAs degrade with time in both untreated and iron-treated cells as seen by accumulation of single-labelled MS2 spots. In order to monitor mRNA degradation, we quantified the number of intact mRNAs as well as stabilized degradation intermediates per cell in a high-throughput manner in hundreds of cells, as described previously (Voigt et al., 2019). Since the quantification of intact and decay fragments is subject to the detection efficiency of the smFISH probes used, the detection efficiency of the smFISH probes used in this study were calculated as described previously (Horvathova et al., 2017) (Figure S4). The fraction of intact mRNAs per cell was then corrected for the detection efficiency of the probes (Figure 2D). Interestingly, the fraction of intact mRNAs per cell reduced faster in iron-treated cells as compared to untreated control cells indicating that mRNAs undergoing translation were also degrading faster.

To determine the kinetics of mRNA decay using our smFISH data, we fit a mathematical model describing the turnover of the intact cytoplasmic transcripts and stabilized decay intermediates to the time evolution of the mean number of RNA species (Horvathova et al., 2017). Interestingly, the resulting degradation rate of cytoplasmic mRNAs in iron-treated cells was faster with a half-life of 3.00±0.75 hours as compared to 5.12±0.86 hours in untreated cells (Figure 2E). In a control TREAT reporter that lacked the IRE, we found that iron had no effect on mRNA stability (supplementary Figure S5). Recently, it has been shown that the ribosome can directly interact with the Ccr4-Not complex, which coordinates mRNA decay through deadenylation and decapping (Buschauer et al, 2020). In order to determine if translating mRNAs were also deadenylated faster than non-translating ones, we measured poly(A) tails in untreated and iron treated cells (Figure S6). Poly(A)-tail shortening was observed over time after induction, which correlated with the decay kinetics of the IRE-TREAT reporter (Figure S6B, S6C). This data indicates that the process of translation (initiation, elongation or termination) can promote mRNA degradation.

### Single-molecule measurements of nonsense-mediated decay kinetics

In order to further validate that our measurements could detect the coupling between translation and mRNA decay, we characterized the translation-dependent degradation of an mRNA that contained a pre-mature termination codon (PTC) and would be subject to the quality control pathway known as nonsense-mediated decay (NMD). During the initial rounds of translation, a PTC is recognized as an aberrant termination event and the transcript is targeted for degradation to prevent accumulation of truncated polypeptides (Kurosaki et al., 2019). To this end, we generated an IRE NMD reporter that contained the well-characterized Triose-Phosphate Isomerase (TPI) in the coding sequence, which contains seven exons and six introns, and was fused to the C-terminus of Renilla luciferase. In our IRE NMD reporter, a PTC was introduced in exon 5 (TPI-PTC-160) whereas an identical reporter without the PTC served as a control (TPI-wildtype) (Hoek et al., 2019)(Figure 3A). These reporters were integrated into a single genomic locus in HeLa cells under the control of a doxycycline-inducible promoter and used to study the kinetics of degradation as described above.

**Figure 3.**
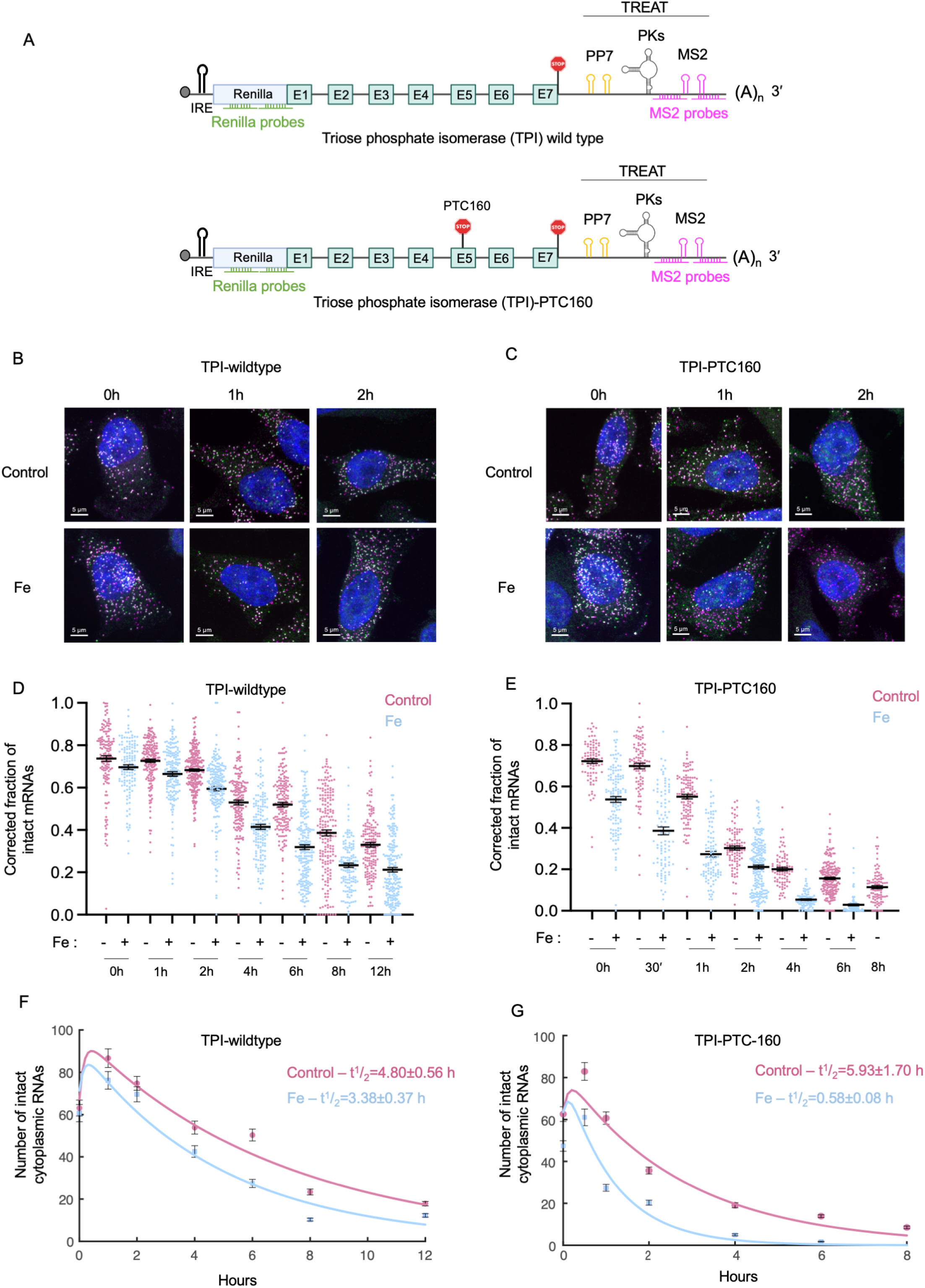
Decay kinetics of NMD reporters using IRE-TREAT. **A.** Schematic representation of reporters to study non-sense mediated decay in presence and absence of translation. The ORF contains 7 exons and 6 introns of the Triose Phosphate Isomerase (TPI) gene. The control reporter is wildtype whereas the NMD reporter contains a premature termination codon (PTC) in exon-5 as indicated (PTC-160). The 5’UTR contains the IRE and the 3’UTR contains the TREAT cassette. The reporters were integrated in HeLa cells under doxycycline-inducible promoter. **B. and C.** Representative smFISH images from cells expressing either TPI-wildtype reporter (B) or TPI-PTC160 reporter (C), in presence or absence of iron, at indicated time-points. Magenta spots indicate signal from MS2 probes and green spots indicate signal from Renilla probes. Dual labelled spots appear white representing intact mRNAs and single labelled magenta spots indicates stabilized intermediates, representing degraded mRNAs. **D.** Quantification of mRNA decay from cells expressing TPI-wildtype-TREAT reporter. Y-axis represents fraction of intact mRNAs corrected for the detection efficiency for the smFISH probes (see supplementary figure S4). Each dot represents fraction of intact mRNAs per cell, data-set from 2 independent experiments. Mean and error bars indicating SEM is represented. Peach dots represent control and blue dots represent iron-treated cells. **E.** Quantification of mRNA decay from cells expressing TPI-PTC-160-TREAT reporter. Y-axis represents fraction of intact mRNAs corrected for the detection efficiency of smFISH probes (see supplementary figure S4). Each dot represents fraction of intact mRNAs per cell, data-set from 2 independent experiments. Mean and error bars indicating SEM are represented. Peach dots represent control and blue dots represent Fe-treated cells. **F. and G.** Quantification of number of cytoplasmic intact mRNAs from the smFISH images of cells expressing TPI-wildtype reporter (F) or TPI-PTC-160 reporter (G) in untreated and iron-treated cells. Mean and error bars representing SEM are indicated. A mathematical model was fitted to the data (solid lines) to obtain the degradation rates of the cytoplasmic RNAs. The half-life with standard deviation is indicated.

Consistent with the results obtained with the IRE-TREAT reporter mRNAs, the TPI-wildtype reporter was also degraded faster in iron-treated cells (half-life of 3.4±0.37 hours) as compared to untreated cells (half-life of 4.8±0.56 hours) (Figure 3B, 3D and 3F). As expected, however, the TPI-PTC-160 reporter mRNAs were rapidly degraded in iron treated as compared to untreated cells (Figure 3C and 3E). About 80% percent of the cytoplasmic TPI-PTC-160 mRNAs were degraded within 2 hours (Figure 3B, 3C), which is similar to previously measurements (Hoek et al., 2019). It should be noted that the TPI-PTC-160 reporter mRNAs also displayed faster decay kinetics in untreated cells, as compared to the IRE-TREAT and TPI-wildtype transcripts, and this is likely due to the basal level of translating mRNAs in untreated cells (Figure 1C, 1E and 1F). The decay kinetics for TPI-PTC-160 mRNAs were modelled with a correction factor to account for fraction of translating mRNAs in untreated and iron-treated cells we observed for IRE-SunTag mRNAs (Figure 3G). In the absence of iron, the corrected half-life of TPI-PTC-160 reporter was calculated to be 5.93±1.70 hours, which is comparable to half-life of TPI-wildtype reporter as well as IRE-TREAT. In iron-treated cells, the TPI-PTC-160 mRNAs degraded rapidly with half-life of 0.58±0.08 hours that is much faster compared to both the TPI-wildtype and IRE-TREAT reporters. Our assay was able to detect the rapid decay of a PTC-containing mRNA, where translation is tightly coupled to decay. Additionally, the difference in the rates of translation-dependent decay we observe for IRE-TREAT and TPI-PTC-160 transcripts indicates the potential of this approach to distinguish different molecular mechanisms.

### miRNAs accelerate degradation independent of translation

miRNAs are *trans*-acting factors that function in complex with Argonaute proteins to recognize their cognate binding sites in the 3’UTRs of transcripts to repress translation as well as cause the degradation of the target mRNAs. Previously, it has been shown that miRNAs associate with translating mRNAs based on the enrichment of miRNAs in polysome fractions (Cottrell et al., 2017; Maroney et al., 2006; Tat et al., 2016). Additionally, lncRNAs, which are not translated, that contained miRNA-binding sites were found to be refractory to miRNA-mediated destabilization (Biasini et al., 2021). In order to determine if translating mRNAs are preferentially degraded by miRNAs, we introduced three let7 miRNA binding sites in the 3’UTR of IRE-TREAT reporter (let7-TREAT) in between the stop-codon and the TREAT cassette (Figure 4A). The let7 miRNA was chosen because it is abundantly expressed in HeLa cells and expression of a reporter with let7 miRNA binding sites has been previously shown to be effectively repressed (Meijer et al., 2013; Pillai et al., 2005). Another reporter with three let7 binding sites in the reverse orientation (let7R-TREAT) was also generated as a control. These reporters were integrated at a single genomic locus in HeLa cells under a doxycycline inducible promoter as described previously.

**Figure 4.**
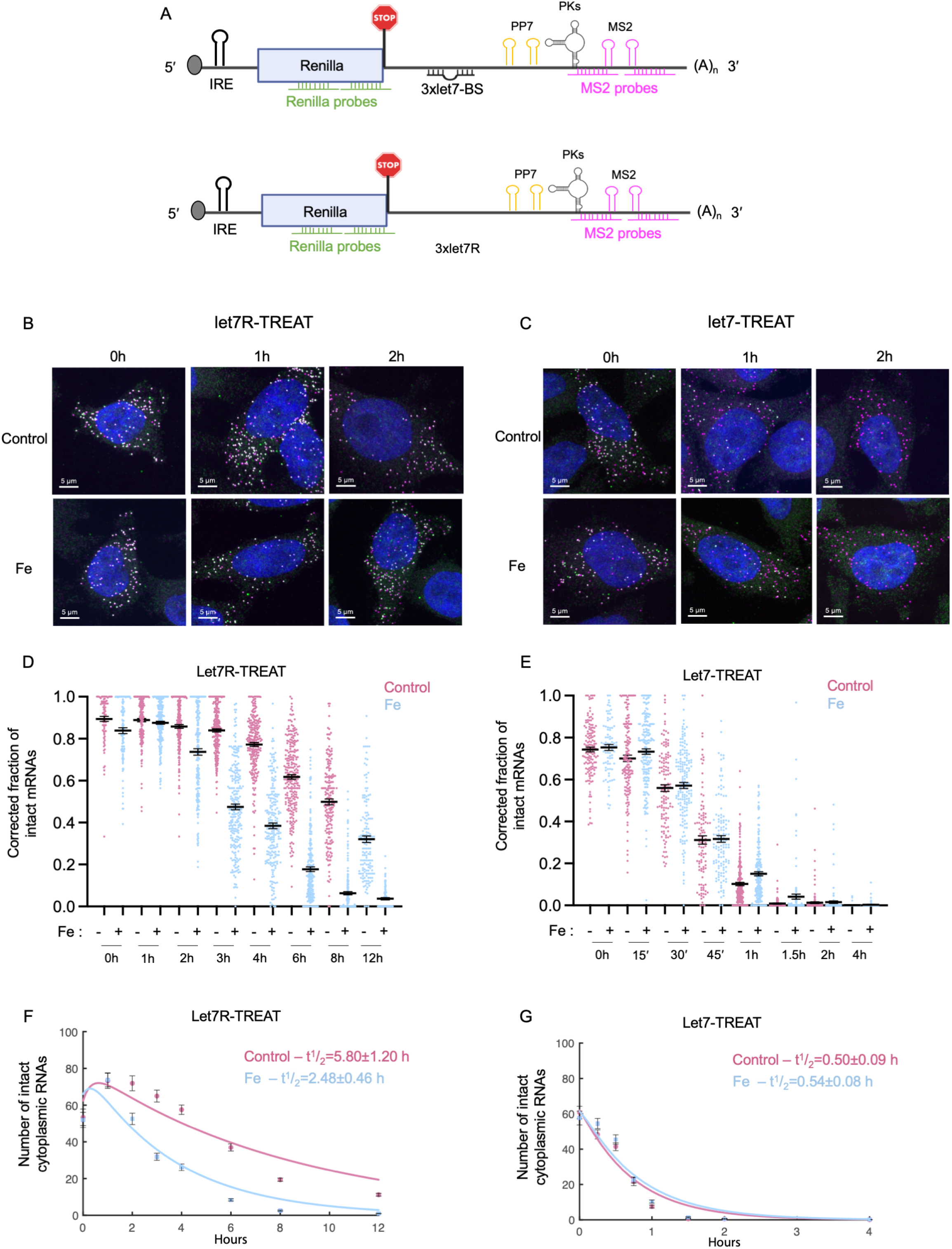
Decay kinetics of miRNA reporters measured using IRE-TREAT. **A.** Schematic representation of miRNA-IRE-TREAT reporters. 3x-let7 miRNA binding site were inserted between the stop codon and the TREAT cassette in the previously describes IRE-TREAT reporter (Figure 2A). Another control reporter was generated with 3x-let7 miRNA binding sites in reverse orientation (3x-let7R). Both the reporters were integrated in HeLa-11ht cells under a doxycycline inducible promoter. smFISH probes targeting Renilla in ORF (upstream PKs) and MS2 stem-loops in 3’UTR (downstream PKs) are indicated. **B. and C.** Representative smFISH images from cells expressing either let7R-IRE-TREAT (B) reporter or let7-IRE-TREAT (C) reporter, in presence or absence of Fe, at indicated time-points. Magenta spots indicate signal from MS2 probes and green spots indicate signal from Renilla probes. Dual labelled spots appear white representing intact mRNAs and single labelled magenta spots indicates stabilized intermediates, representing degraded mRNAs. **D.** Quantification of mRNA decay from the smFISH images of cells expressing let7R-IRE-TREAT. Y-axis represents fraction of intact mRNAs corrected for the detection efficiency of smFISH probes. Each dot represents fraction of intact mRNAs per cell, data-set from 2 independent experiments. Mean and error bars indicating SEM are represented. Peach dots represent control and blue dots represent Fe treated cells. **E.** Quantification of mRNA decay from the smFISH images of cells expressing let7-IRE-TREAT. Y-axis represents fraction of intact mRNAs corrected for the detection efficiency for the smFISH probes. Each dot represents fraction of intact mRNAs per cell, data-set from 2 independent experiments. Mean and error bars indicating SEM is represented. Peach dots represent control and blue dots represent Fe treated cells. **F. and G.** Quantification of number of cytoplasmic intact RNAs from the smFISH images of cells expressing let7R-IRE-TREAT (F) or let7-IRE-TREAT (G) in untreated and iron-treated cells. Mean and error bars representing SEM is indicated. A mathematical model was fitted to the data (solid lines) to obtain the degradation rates of the cytoplasmic RNAs. The half-life with standard deviation is indicated.

We first measured the degradation rate of the let7R-TREAT reporter in the presence and absence of iron and found that its degradation was similar to both the IRE-TREAT and TPI-wildtype transcripts in both conditions (Figure 4B and 4D). Specifically, in iron-treated cells the half-life was 2.48±0.46 hours and in untreated cell the half-life was 5.80±1.20 hours (Figure 4F). Interestingly, the let7-TREAT reporter was rapidly degraded in both untreated as well as iron-treated cells (Figure 4C and 4E), as shown by single-labelled MS2 spots whereas the majority of the control let7R-TREAT mRNAs remained intact by 2 hours (Figure 4B). Quantification of the images revealed that about 85% percent of let7-TREAT mRNAs were degraded within one hour in both untreated as well as iron-treated cells (Figure 4E). The smFISH data was used to model the decay kinetics and it was found that the degradation rates of cytoplasmic let7-TREAT mRNAs were identical in both untreated (half-life of 0.50±0.08 hours) and iron-treated (half-life of 0.54±0.08 hours) cells (Figure 4G). Consistent with the rapid degradation of let7-TREAT transcripts, these mRNAs were also found to be deadenylated at the earliest time points (Figure S7), which is similar to previous measurements of miRNA-mediated deadenylation (Chen et al., 2009; Eisen et al., 2020a). These results indicates that recruitment of miRNAs to target mRNAs and their subsequent degradation is independent of its translation status.

### miRNAs predominantly function to degrade target mRNAs

While our let7-TREAT data indicates that miRNAs can target transcripts for degradation regardless of their translation status, these experiments could not determine if repression of translation occurred before degradation. To simultaneously monitor the translation and degradation of a single mRNAs, we modified the let7-TREAT and let7R-TREAT reporters to contain the SunTag cassette in the coding sequence (Figure 5A). We used a smFISH-IF protocol to quantify translation in fixed cells by immunofluorescence using an anti-GFP antibody that enhances the signal of the scFv-GFP bound to the SunTag epitopes and mRNA decay was measured by smFISH using probes targeted to Renilla and the MS2 stem-loops (Figure 5A). In this experiment, it is possible to detect three distinct populations of mRNAs based on their labeling; a) triple labelled spots indicating intact mRNAs undergoing translation, b) dual labelled (MS2+Renilla) spots indicating non-translating intact mRNAs and c) single labelled MS2 spots indicating stabilized degraded intermediates (Figure 5B). Consistent with our previous live cell experiments using the IRE-SunTag reporter, we detected a higher fraction of translating mRNAs, indicated by triple labelled spots, in iron-treated cells than in untreated control cells (Figure 5B). Importantly, we also do not detect SunTag signal on MS2-only spots indicating the specificity of these measurements.

**Figure 5.**
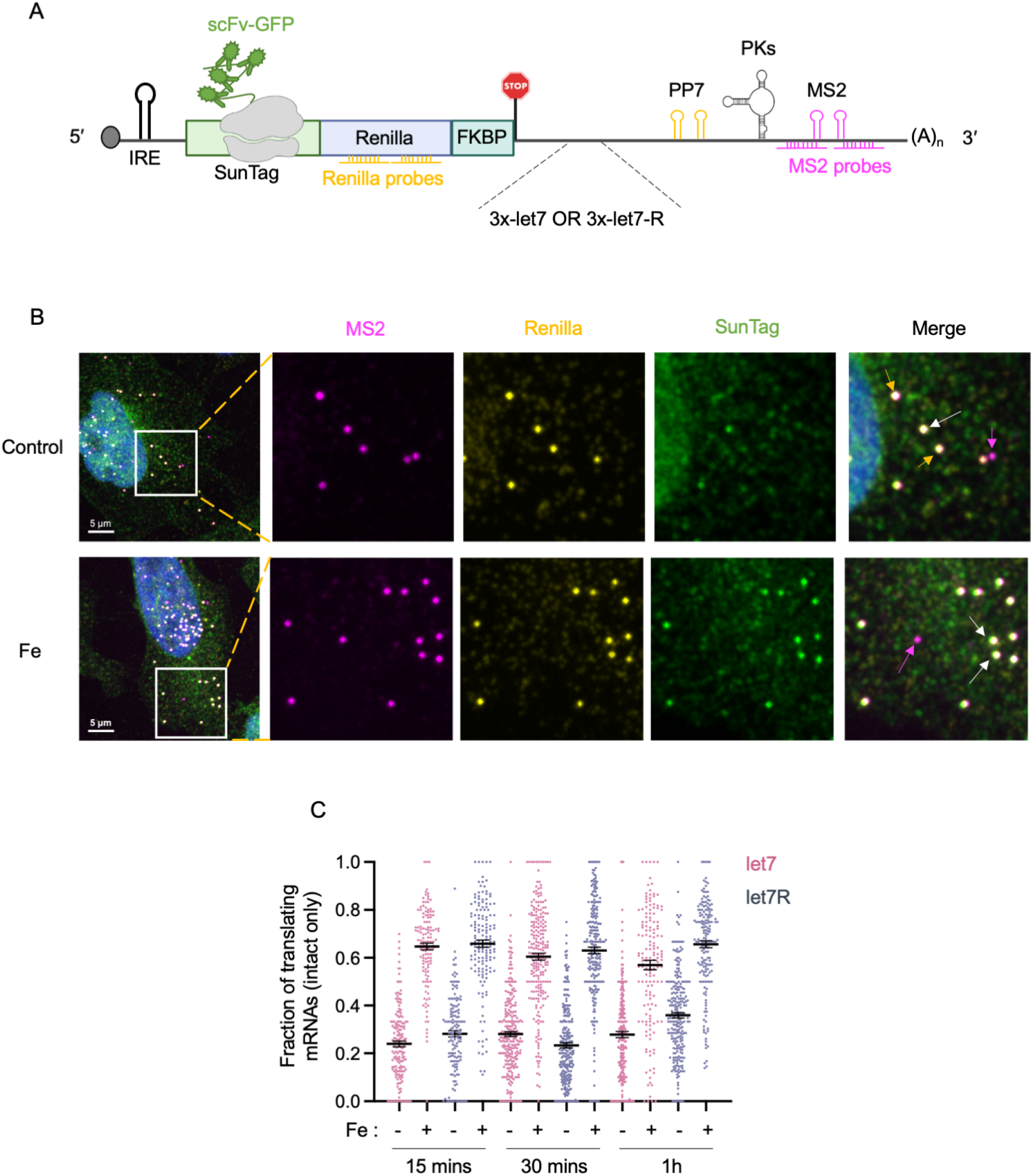
Three-color single-molecule imaging of translation and RNA decay to study miRNA function. **A.** Schematic representation of miRNA-IRE-SunTag-TREAT reporters used for 3-color smFISH-IF. The SunTag cassette was inserted in the ORF of let7-IRE-TREAT and let-7R-IRE-TREAT reporters. Both the reporters were integrated in HeLa-11ht cells expressing scFV-GFP. smFISH probes targeting Renilla in ORF (upstream PKs) and MS2 stem-loops in 3’UTR (downstream PKs) are indicated, while anti-GFP antibody was used to detect the SunTag signal on mRNAs. **B.** Representative image of smFISH-IF in presence and absence of Fe. Magenta spots indicate signal from MS2 probes and yellow spots indicate signal from Renilla probes. Green spots represent SunTag signal from translating mRNAs. The magenta arrow indicates stabilized intermediate representing degraded mRNA. White arrow indicates intact mRNA undergoing translation and yellow arrow indicates non-translating intact mRNA. **C.** Quantification of fraction of translating mRNAs (from intact mRNAs only) per cell from smFISH-IF images of cells expressing either let7-SunTag-TREAT reporters (peach spots) or let7R-SunTag-TREAT (blue spots), at indicated time-points, in untreated or iron-treated cells. Each dot on the plot corresponds to data from one cell. Mean and SEM are indicated.

We then compared the translation and decay of let7-SunTag-TREAT and let7R-SunTag-TREAT mRNAs. As expected, the let7-SunTag-TREAT transcripts were found to degraded faster than the let7R-SunTag-TREAT mRNA, albeit with slightly slower kinetics than the let7-TREAT reporter without the SunTag (Figure S8). Interestingly, at all timepoints the fraction of translating mRNAs remained similar for both let7-SunTag-TREAT and let7R-SunTag-TREAT reporters in both the presence and absence of iron (Figure 5C). As miRNA-mediated decay was already detected at 1h (Figure S8), these experiments indicated that let7 miRNA was unable to strongly repress translation prior to (or independently of) decay.

In our fixed cell smFISH-IF experiments, we only quantified a transcript as translating or not translating and, therefore, could not determine if the let7 miRNA reduced ribosome occupancy prior to degradation. In order to test if there was a difference in the translation efficiency of let7-SunTag-TREAT transcripts compared to let7R-SunTag-TREAT, live-cell imaging of translation sites was performed (Figure 6A, S9A and S9B). Consistent with our previous IRE-SunTag experiments, the SunTag intensity of both let7-SunTag-TREAT and let7R-SunTag-TREAT mRNAs increased in iron-treated cells (Figure 6B). The fraction of translating mRNAs for let7-SunTag-TREAT reporters was slightly lower, but statistically insignificant, than the control reporters in both untreated and iron-treated cells (Figure 6C). Analysis of SunTag intensities also revealed a similar distribution of ribosomes on let7-SunTag-TREAT transcripts as compared to let7R-SunTag-TREAT mRNAs (Figure 6D, S9C). Taken together, our results indicate that let7 miRNA-mediated regulation of mRNAs occurs primarily via mRNA degradation and that inhibition of translation has a modest, if any, role.

**Figure 6.**
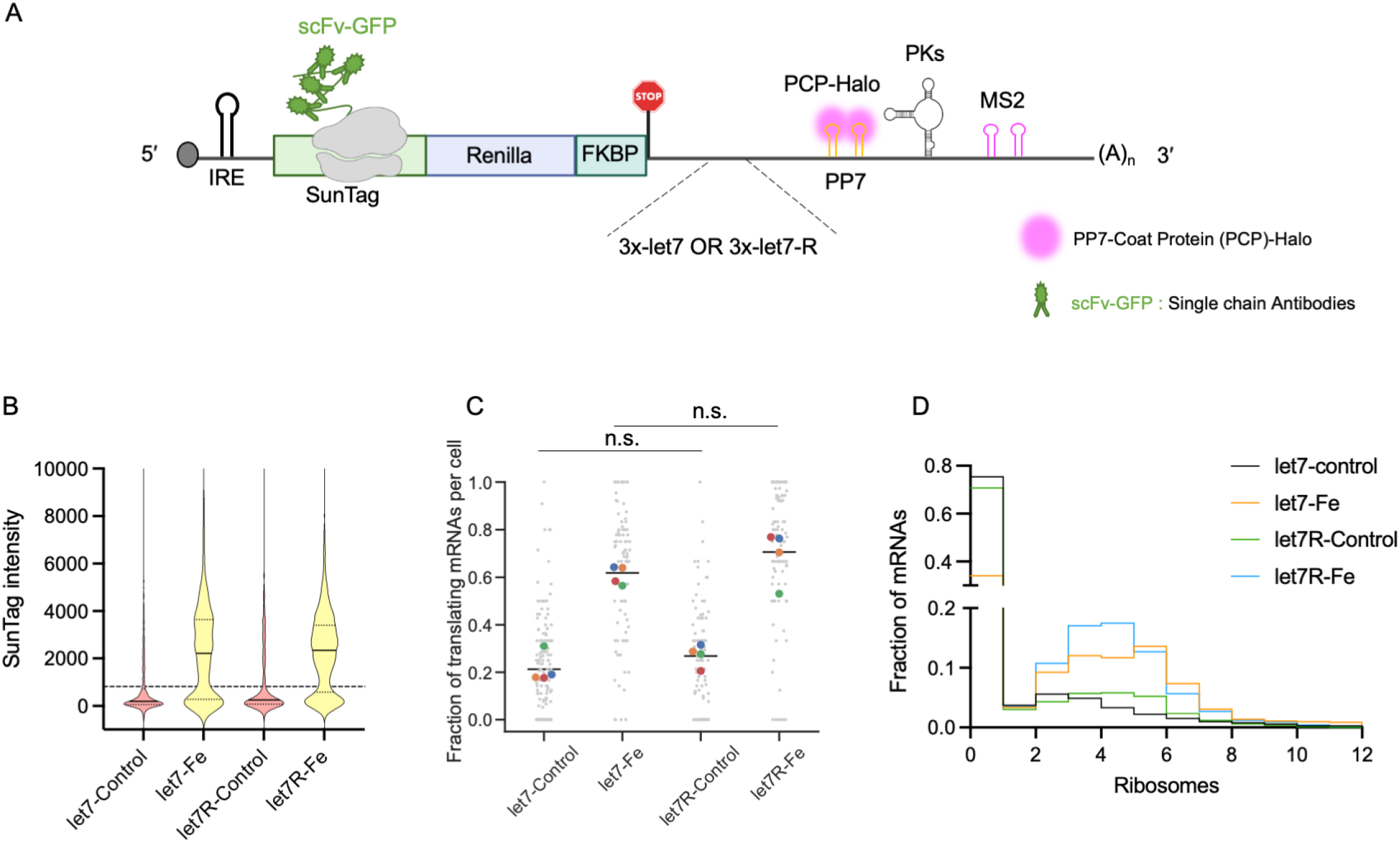
Live-cell imaging of let7-SunTag-TREAT reporters and let7R-SunTag-TREAT reporters. **A.** Schematic showing the strategy used for live-cell imaging. The reporter RNAs (let7-SunTag-TREAT or let7R-SunTag-TREAT) were expressed in HeLa cells stably expressing scFV-GFP and PP7-coat protein fused to HaloTag. **B.** Quantification of the SunTag intensities corresponding to the SunTag-let7-TREAT mRNAs in untreated cells (1358 mRNAs) or iron-treated cells (952 mRNAs) and for the SunTag-let7R-TREAT mRNAs in untreated cells (1026 mRNAs) or iron-treated cells (1195 mRNAs) from 4 independent experiments. The background GFP intensity of the cell was subtracted from the measured SunTag intensity of each mRNA. The dotted line represents the cut-off based on intensity of a single SunTag polypeptide. **C.** The distribution of fraction of translating mRNAs per cell. let7-SunTag-TREAT-control (153 cells), let7-SunTag-TREAT-Fe (88 cells), let7R-SunTag-TREAT-control (102 cells) and let7R-SunTag-TREAT-Fe (95 cells) from 4 independent experiments. n.s. indicates not significant. Color-coded dots represent averages from individual experiment set. **D.** Quantification of number of ribosomes on mRNAs represented in figure 6B. Y-axis represents relative fraction of mRNAs from 4 independent experiments. The frequency of mRNAs with number of ribosomes for all the mRNAs represented.

## DISCUSSION

In this study, we combined transcript-specific control of translation with singlemolecule imaging of translation and mRNA decay to investigate the coupling between these steps in the gene expression pathway. By directly measuring the translation status and degradation of individual mRNAs, we find that the process of translation promotes mRNA decay. While this provides a simple mechanism for the coupling between the transcriptome and the proteome, it also provides a reference point for deciphering the mechanisms that result in discordant mRNA and protein levels. In this context, it is interesting to reexamine why highly abundant transcripts (e.g. housekeeping genes) that have high translation efficiencies are also some of the most stable transcripts (Eraslan et al., 2019; Vogel et al., 2010). If translation does indeed promote mRNA decay, these transcripts would require mechanisms to counteract this effect. In the case of 5’TOP (terminal oligopyrmidine) transcripts, which encode ribosomal proteins and translation factors, the *trans*-acting factor Larp1 has been implicated in both regulating translation and mRNA stability (Berman et al., 2021). If other housekeeping genes employ similar strategies will be an important area of future research.

While our data indicates that translation promotes mRNA decay, the underlying mechanism is not fully understood. We find that translation results in rapid deadenylation, which is consistent with a recent study that measured the dynamics of poly(A)-tail length and found that deadenylation rates correlate with the mRNA halflife (Eisen et al., 2020b). In principle, however, the increased deadenylation rates that we observe upon translation could arise from a process occurring during initiation, elongation or termination. One possibility is that the interaction between the ribosome and the Ccr4-Not complex that has been shown to connect codon optimality with mRNA stability, may play a more general role in promoting the decay of translating versus non-translating mRNAs (Buschauer et al., 2020). Alternatively, recently it has been shown that ribosome readthrough of stop codons occurs more frequently than anticipated and could also results in mRNA destabilization (Boersma et al., 2019).

miRNAs are well known to post-transcriptionally repress mRNAs either by inhibiting translation, accelerating mRNA decay or a combination of both (Bartel, 2018). While this manuscript was in preparation, two other studies using single-molecule translation site imaging characterized the effect of miRNAs on repression of gene expression (Cialek et al., 2021; Kobayashi and Singer, 2021). While all three studies agree that miRNA-mediated targeting of transcripts occurs rapidly after export to the cytoplasm, the other studies suggest that translational inhibition occurs prior to mRNA decay and that translating mRNAs are preferentially targeted by miRNAs. Our results, however, indicate that miRNAs promote mRNA decay independent of the translation status of the transcript and that inhibition of translation does not precede degradation. It is to be noted that, while all three studies utilize single-molecule translation-site imaging, we combine it with TREAT which allowed us to also directly visualize mRNA decay, together with translation. While we cannot exclude the possibility that the specific cell lines or miRNAs investigated in the different studies resulted in these different observations, we believe this highlights the necessity of direct single-molecule measurements of mRNA decay in order to correctly determine its contribution to repression.

Finally, the successful development of mRNA vaccines targeting SARS-CoV-2 has ushered in new era in drug discovery (Jackson et al., 2020). As the amount of protein generated by a therapeutic mRNA may become a critical parameter for the success of drug or vaccine, it will become increasingly important to understand the connection between translation and mRNA decay. Since our results indicate that translation promotes mRNA decay, strategies that actively stabilize therapeutic mRNAs against this effect maybe beneficial.

## ACKNOWLEDGMENTS

This work was supported by the Novartis Research Foundation (J.A.C), the Swiss National Science Foundation grant 31003A_182314 (J.A.C), the SNF-NCCR RNA& Disease (J.A.C), an EMBO fellowship ALTF 1203-2018 (D.M) and an EMBO fellowship ALTF-598-2019 (P.D). This project has received funding from the European Union’s Horizon 2020 research and innovation programme under the Marie Skłodowska-Curie Grant Agreement No. 843212. The authors thank members of the Chao lab for helpful discussions and critical reading of the manuscript, L. Gelman and S. Bourke for microscopy support and H. Kohler for cell sorting. We also thank H. Grosshans and W. Fillipowicz for helpful feedback on the manuscript. T. Hochstoeger and F. Voigt from the Chao lab are thanked for PCP-Halo plasmid and virus.

## AUTHOR CONTRIBUTIONS

P.D. performed all experiments and image analysis. E.G. performed poly(A)-tail measurements. G.R. helped with calculating mRNA decay rates. D.M. developed the workflows for analysis of live-cell imaging data. P.D. and J.A.C. wrote the article with input from all the authors.

## METHODS

### Cell culture

Previously described HeLa-11ht cells were used for all the experiments in this study (Weidenfeld et al., 2009). The cell line contains a Flp-RMCE site (recombinase mediated cassette exchange), which allows single-copy integration of the reporters used in the study. For doxycycline induction of transcription, the cells also express reverse-tetracycline controlled transactivator (rtTA2S-M2). HeLa cells were cultured in DMEM containing 4.5 g/L glucose, antibiotics penicillin and streptomycin (100 μg/ml), 4 mM L-glutamine and 10% fetal bovine serum (FBS) and were maintained in 37°C and 5% CO_2_. Lipofectamine 2000 (Invitrogen) was used for transfections as per manufacturer’s instructions and transfections were performed with Opti-MEM (Thermo Fisher Scientific).

### Cell lines and plasmids

Different reporter plasmids generated in this study are described below. Cell lines expressing the individual reporter were generated as described previously (Weidenfeld et al., 2009). Briefly, cells were seeded in 6-well plate and were transfected with 1 μg of plasmid expressing FLPe recombinase (addgene plasmid: 20733) (Beard et al., 2006) along with 1 μg of either reporter plasmid described below. The reporter genes are flanked by Flp-recombinase target sites, allowing recognition by the FLPe recombinase protein. 24 hours post transfection, the cells were treated with puromycin (5 mg/ml) (Invivogen) for two days to select for transfected cells. Cells were selected for successful integrators by maintaining them in 50 mM glanciclovir for 10 days. The cell line integrated with the SunTag-Renilla reporter together with scFV-GFP and NLS-stdMCP-std-Halo was described previously (Wilbertz et al., 2019). IRE-TREAT, IRE-TPI-wildtype-TREAT, IRE-TPI-PTC-160-TREAT, let7-IRE-TREAT and let7R-IRE-TREAT reporters were integrated in the HeLa-11ht cell line. The IRE-SunTag reporter was integrated in previously established HeLa cell line stably expressing scFv-GFP (Addgene plasmid: 104998) and NLS-stdMCP-stdHalo fusion protein (Addgene plasmid: 104999) (Voigt et al., 2017). Let7-SunTag-TREAT and let7R-SunTag-TREAT were integrated in HeLa-11ht cell line expressing scFv-GFP and NLS-PCP-Halo fusion protein.

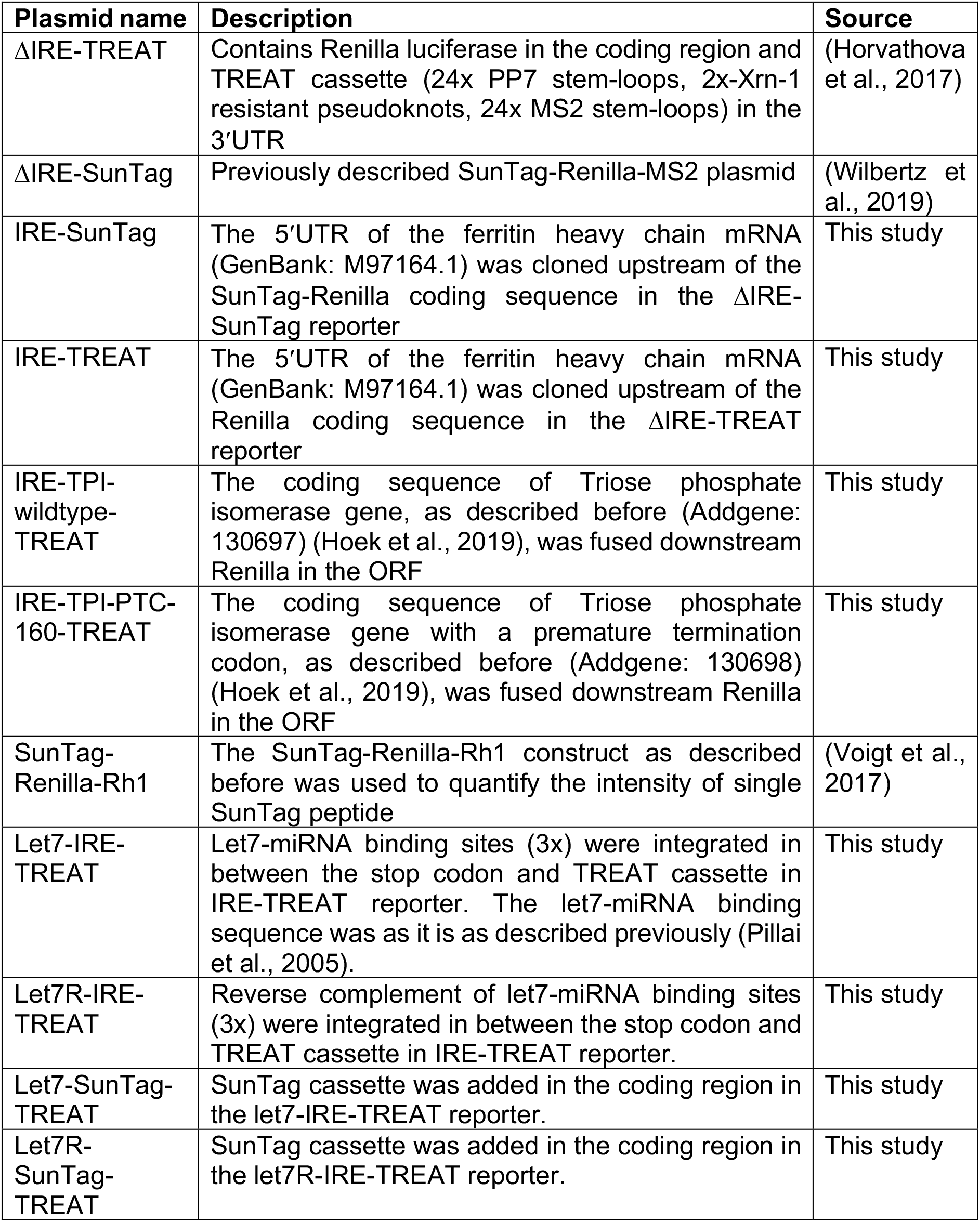

### Single-molecule Fluorescence in situ hybridization (smFISH) and immunofluorescence (IF)

HeLa cells were seeded on glass coverslips (high precision, 170 μm thick, 18 mm diameter, Paul Marienfeld GmbH) placed in a 12-well tissue culture plate (0.1×10^6^ cells per well) and grown for 48 hr. In order to activate the translation of the reporter, 150 μM of ferrous ammonium citrate was added to the culture medium 1h before the induction of transcription and was maintained throughout the experiment. The smFISH protocol used to study the decay kinetics of all the TREAT reporters used in the study was performed as described previously (Horvathova et al., 2017). At indicated time points, cells were fixed in 4% paraformaldehyde in PBS for 10 minutes and subsequently permeabilized by overnight incubation in 70% ethanol at 4°C. Cells were pre-blocked in wash buffer (2xSSC (Invitrogen), 10% v/v formamide (Ambion)) for 15 minutes. The cells were hybridized in hybridization buffer (150 nM of smFISH probes, each targeting Renilla and MS2 stem-loops, 2xSSC, 10% v/v formamide, 10% w/v dextran sulphate (Sigma)) for 4 hours at 37°C and then washed twice with wash buffer for 30 minutes each. The coverslips were mounted on slides in ProLong Gold antifade containing DAPI (Molecular Probes). 20-nucleotide long oligos targeting MS2 stemloops and Renilla luciferase were designed using Stellaris probe designer and the oligos were conjugated with Atto-565 and Atto-633 dyes respectively, as previously described (Gaspar et al., 2017).

For visualizing the SunTag signal, smFISH-IF was performed after fixing the cells at indicated timepoints in 4% paraformaldehyde. The cells were permeabilized in 0.5% Triton-X solution for 10 minutes. The cells were preblocked in wash buffer, described above, also containing 3% BSA (Sigma) for 30 minutes at room temperature. The cells were washed twice in 1x PBS and hybridized in the aforementioned hybridization buffer containing 1:1000 diluted anti-GFP antibody (Aves labs GFP-1010) for 4 hours at 37°C. After hybridization, cells were washed with wash buffer for 30 minutes and incubated in anti-chicken IgY secondary antibody conjugated with Alexa-fluor 488 for 30 minutes. The coverslips were washed twice in 1x PBS and were mounted on glass slides in ProLong Gold antifade containing DAPI (Molecular Probes).

Images were acquired using a Zeiss AxioObserver7 inverted microscope equipped with a Yokogawa CSU W1-T2 spinning disk confocal scanning unit, a Plan-APOCHROMAT 100x 1.4 NA oil objective, a sCMOS camera and an X-Cite 120 EXFO metal halide light source. 21 z-stacks were performed in 0.24 μm steps. The exposure times were 800 ms for Atto-633, 500 ms for Atto-565 and 50 ms for the DAPI channel.

### Analysis of smFISH and smFISH/IF data

Detection and quantification of the single mRNA spots in smFISH images was performed in KNIME as described previously (Voigt et al., 2019). Maximum intensity projections of the images were used for spot detection using the “spot detection” node in KNIME based on intensity thresholds, that were kept consistent for the all timepoints in a given experiment for each channel. The segmentation of nucleus was performed based on Otsu thresholding method (Otsu, 1979) in KNIME. The background nonspecific intensity from the smFISH probes was used to segment the cytoplasm. The detected spots in each channel were co-localized using nearest neighbor search in different spot channels and 2 spots within maximum distance of 3 pixels (321 nm) were considered as co-localized.

In order to correct for the efficiency of probes to detect the mRNAs in smFISH protocols, the detection efficiency of the probes was calculated as described earlier (Horvathova et al., 2017). A reporter RNA lacking the Xrn-1 resistant pseudoknots was expressed in cells and smFISH was performed using MS2 and Renilla probes (Figure S4A, S4B). Since these reporters would not lead to generation of stabilized decay intermediates, all mRNAs should be intact and dual labelled. Based on the number of single labelled spots, the detection efficiency was calculated and represented in figure S4B.

### Modeling of mRNA half-lives from smFISH measurements

In the absence of transcription, we assumed that the export of the intact nuclear mRNA into the cytoplasm, the degradation of intact cytoplasmic mRNA into stabilized 3; ends and the degradation of the stabilized 3’ends are described by three single-step Poisson processes with rates α, β and γ, respectively. We assume that only the degradation rate of the intact cytoplasmic mRNA differs between control and iron treatment (i.e. *β_c_* for control and *β_i_* for iron treatment). Hence, the mean turnover of these RNA species is described by the following three rate equations:

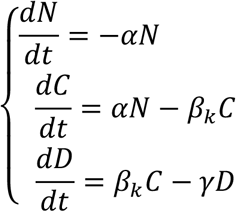

where *N* is the number of intact nuclear mRNA, *C* is the number of cytoplasmic mRNAs, and *D* is the number of stabilized 3’ ends, respectively, and *k* = *c* for the control and *k* = *i* for the iron treatment.

We first fitted the solutions of this model to the control population-averaged intact nuclear mRNA, cytoplasmic mRNAs, and the stabilized 3’ ends FISH data. We obtained the best fit parameters *α*^*^, 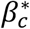 and *γ*^*^. Then we fitted the solutions of the model to the iron treated population-averaged intact nuclear mRNA, cytoplasmic mRNAs, and the stabilized 3’ ends FISH data, fixing the parameters *α* and *γ* to the values of the control best fit values *α*^*^, *γ*^*^, respectively, and letting the degradation rate *ßι* free. All the fits were calculated with the *Matlab* function *lsqcurvefit*.

### Modeling of Nonsense-mediated decay (NMD) of mRNAs

Since NMD transcripts are well established to undergo translation-dependent decay, we assumed that intact cytoplasmic mRNAs degrade at rate *β_t_* when they are translating and at rate *β_nt_* when they are not translating. Based on experiment measurements (Figure 4E), we assumed a fixed fraction of translating mRNAs which is equal to 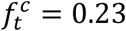 for the control 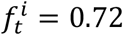 in the iron treatment. Hence the mean turnover of these RNA species is described by the following three rate equations:

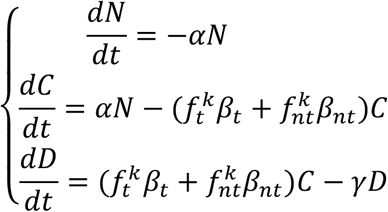

where *k* = *c* for the control and *k* = *i* for the iron treatment.

We fitted simultaneously the solutions of this model to the control population-averaged FISH data and to the iron treated population-averaged FISH data. We obtained the best fit parameter values for *α, β_t_, β_nt_* and *γ*. All the fits were calculated with the *Matlab* function *lsqcurvefit*.

### Live cell Imaging

Cells were grown on 35mm glass-bottom dish (ibidi GmbH) for two days prior to imaging. In order to label the endogenous MCP-Halo proteins, the cells were supplemented with 100 nM of JF549 HaloTag ligand, obtained from L. Lavis (Janelia Research Campus) (Grimm et al., 2015), for 20 minutes, after which the medium was replaced and cells were washed in PBS. To control the translation, 150 μM of ferrous ammonium citrate (Sigma, 22896-6) or iron chelator IV-21H7 (Sigma, 681679-M) was supplemented to the culture medium for indicated period of time. To induce the transcription of reporter mRNAs, the growth medium was replaced with a fresh medium containing 1 mg/mL doxycycline. During the course of imaging, cells were maintained in FluoroBrite DMEM (Thermo Fisher Scientific) containing 10% FBS and 4 mM L-glutamine (also containing 150 μM of ferrous ammonium citrate when indicated). Live cell imaging was performed on an inverted microscope Ti2-E Eclipse (Nikon) equipped with CSU-W1 Confocal Scanner Unit (Yokogawa), 2 back-illuminated EMCCD cameras iXon-Ultra-888 (Andor), MS-2000 motorized stage (Applied Scientific Instrumentation) and VisiView^®^ imaging software (Visitron Systems GmbH). The cells were maintained at 37°C in presence of 5% CO_2_ during entire imaging session. To illuminate the cells, 561 Cobolt Jive (Cobolt) and 488 iBeam Smart lasers and VS-Homogenizer (Visitron Systems GmbH) were used. CFI Plan Apochromat Lambda 100X Oil/1.45 objective (Nikon) was used, resulting in a pixel size of 130nm. Both channels were imaged simultaneously and the emissions were acquired in two separate cameras. The camera misalignment was corrected during analysis using the images acquired from imaging 0.5μm fluorescent beads, as described in “Analysis of live-cell single-molecule imaging data” section.

### Analysis of live-cell single-molecule imaging data

To correct for camera misalignment and chromatic aberrations, images of fluorescent TetraSpeck™ beads were used to determine the channel shift (affine transformation) using the Descriptor-based registration plugin (Preibisch et al., 2010) in Fiji (Rueden et al., 2017; Schindelin et al., 2012). Fiji macro was then used to re-apply the transformation model to the movies acquired on the same imaging session.

Single-particle tracking of mRNA and subsequent analysis of translation sites was performed using custom-built workflows in KNIME (Berthold et al. 2009; Dietz et al. 2020). First, we imported the registered 30-frame movies. Individual cells were then manually annotated using the “Interactive Annotator” node in the MCP channel to create regions of interest (ROIs) that include the cytosol without nucleus. Singleparticle tracking was then performed within these ROIs to detect and track spots in the MCP channel using the “Track Spots (Subpixel localization, multi-channel)” node (Eglinger 2019). This node runs TrackMate with the multi-channel intensity analyzer extension (Tinevez et al., 2017), which enabled the measurement of the SunTag fluorescence intensity in the detected MCP foci. Mean track intensity was then calculated from all frames in the mRNA trajectory, and tracks shorter than 3 frames were filtered out. To correct for cell-to-cell variability in the scFv-GFP expression levels, we further subtracted the background fluorescence intensity in the cytosol (defined as mean intensity in the annotated ROI minus the detected trajectories). The background-subtracted SunTag intensities were plotted in GraphPad Prism (Basham, 1997).

To estimate the fraction of actively translating mRNAs and the number of ribosomes per mRNA, we first measured the fluorescence intensity of a single mature SunTag peptide, as described previously (Voigt et al., 2017). This was achieved by imaging the cells expressing Renilla-SunTag-Rh1 fusion proteins which stably associate with actin filaments. The cells were pretreated with 100 μg/ml of puromycin to eliminate the signal from translating mRNAs. The movies of the actin-anchored SunTag peptides were imported into KNIME and similar analysis was performed as described above. In this case, cells with actin-anchored peptides were annotated in the SunTag channel, single-particle tracking was performed in the SunTag channel (since the fully translated peptides do not colocalize with MCP foci), and tracks shorter than 10 frames were filtered out (since the actin-anchored foci are more stationery). We defined the intensity of a mature SunTag peptide as the median intensity of all the track intensities. To calculate the fraction of translating and non-translating mRNAs from the SunTag intensities on mRNA tracks (described in the first paragraph), the intensity of a mature SunTag peptide was used as a cutoff. The fractions of translating mRNAs were plotted in Python (Waskom et al., 2021).

To further convert the SunTag intensities into the number of translating ribosomes per mRNA, we calculated the fluorescence intensity of an average translating ribosome. Since not all ribosomes are associated with a fully synthesized SunTag array, the intensity of the mature SunTag peptide was multiplied by a coefficient (0.71) to correct for the position of ribosomes on the mRNA. This correction assumes a uniform distribution of ribosomes on the mRNA. Only the ribosomes downstream of the SunTag array will be associated with a fully synthesized SunTag array and thus have the fluorescence intensity of the mature SunTag peptide. Ribosomes that occupy the first part of the coding sequence (SunTag array) will have on average half the signal of the fully synthesized SunTag peptide. Using this corrected intensity of an average translating ribosome (and its multiples), we binned the SunTag intensities on mRNA tracks (described in the first paragraph) in order to estimate the number of translating ribosomes per mRNA. As a cutoff between 0 and 1 ribosomes, we used the intensity of 1 average ribosome, in order to better distinguish between weakly translated mRNAs and non-translating mRNAs without a real SunTag signal. The numbers of translating ribosomes per mRNA were plotted as relative fractions using Prism.

## Supplementary Figure Titles and Legends

**Figure S1.**
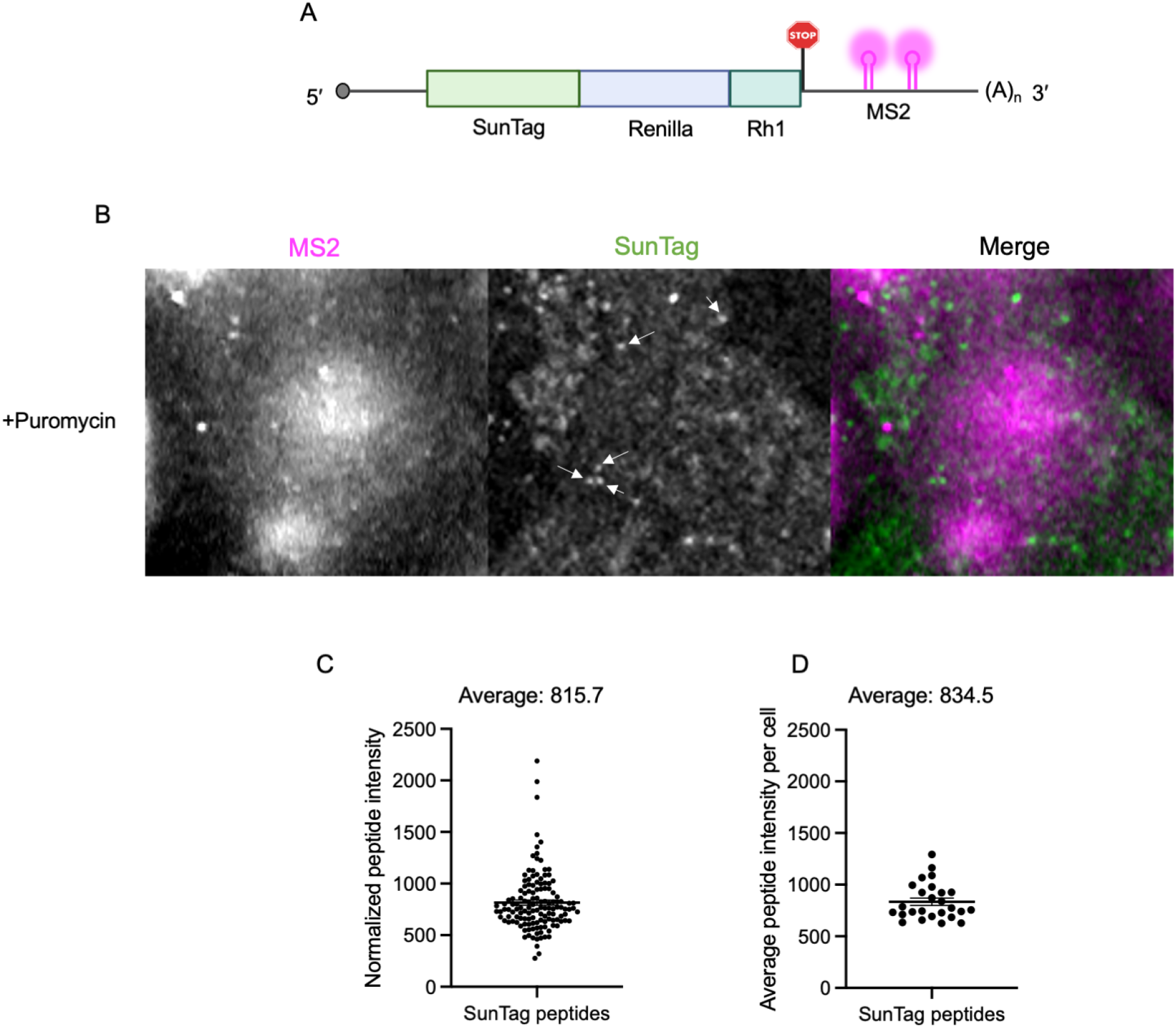
Quantification of intensity of single SunTag polypeptide. **A.** Schematic representation of reporter used to quantify the intensity of single SunTag-polypeptide. Upon translation the mature SunTag polypeptide containing the Rh1 domain associates with Actin-filaments allowing stable imaging of actin-anchored individual SunTag-polypeptides. **B.** Representative image of SunTag-Rh1 reporter expressed in puromycin-treated HeLa cells stably expressing MCP-Halo and scFV-GFP. Imaging was carried out after treating cells with 100 μg/mL puromycin for 10 mins to dissociate ribosomes from the mRNAs. The white arrows indicate individual SunTag-polypeptides bound to actin-filaments (See supplementary video 7). **C.** Quantification of intensities of individual mature SunTag-polypeptides (background subtracted) (134 peptides, 26 cells). Mean±SEM – 815.7±24.45. **D.** Average intensity of mature SunTag-peptides per cell. Mean±SEM – 834.6±34.76.

**Figure S2.**
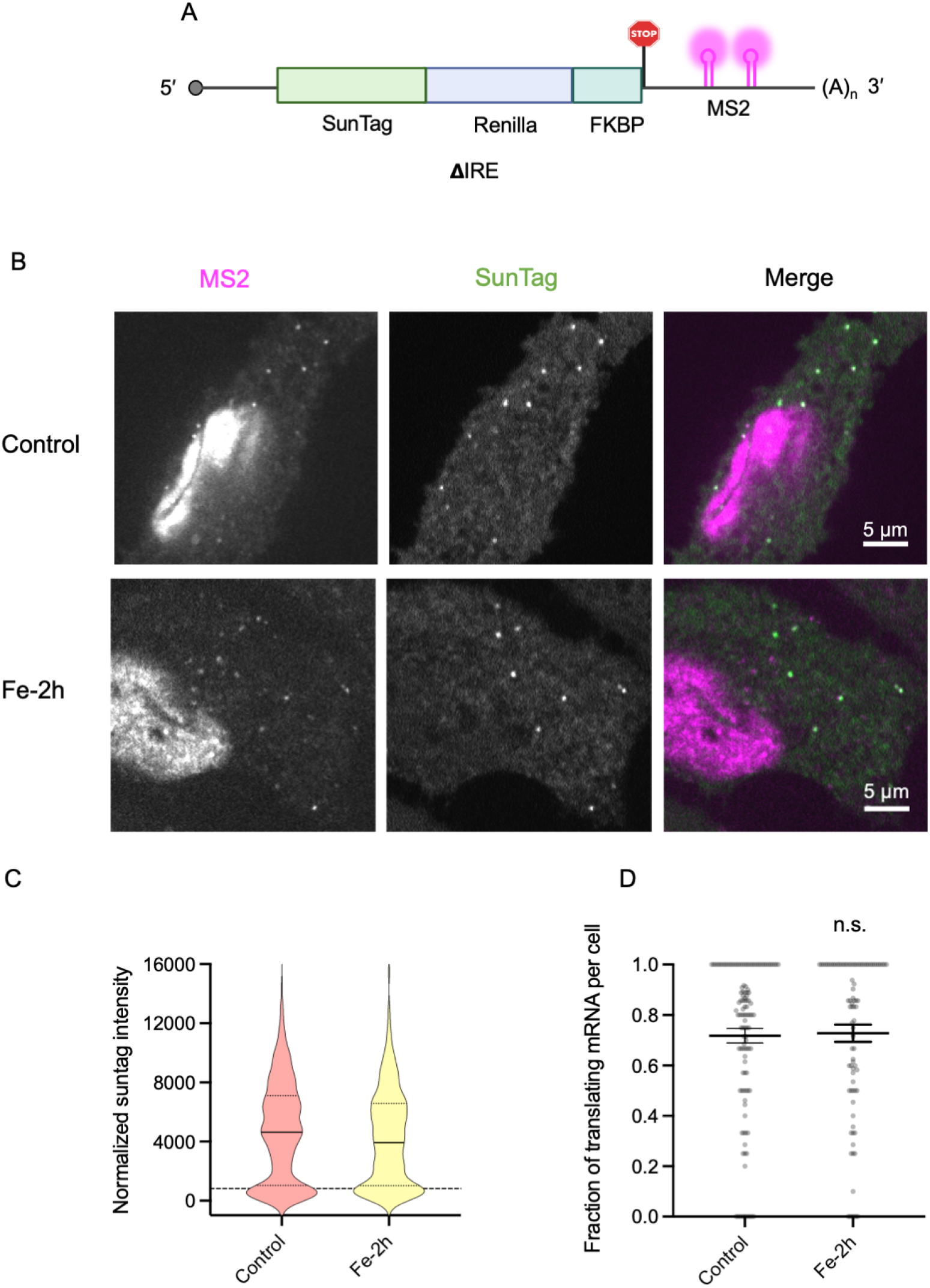
Iron-treatment does not affect the translation of ΔIRE-SunTag reporter. **A.** Schematic representation of the ΔIRE-SunTag reporter used to study translation upon treatment of cells with iron. **B.** Live-cell single-molecule imaging of ΔIRE reporters upon treatment of cells with iron for 2h (Fe-2h) or in absence of Fe (Control). Magenta spots represent mRNAs and green spots represents SunTag indicating translation. **C.** Quantification of the background subtracted SunTag intensities corresponding to the ΔIRE reporter mRNAs in absence of Fe (Control-1092 mRNAs) or in presence of Fe treated for 2h (Fe-2h-758 mRNAs), from 2 independent experiments. The dotted line represents the cut-off based on intensity of single SunTag polypeptide. **D.** Fraction of translating mRNAs per cell in control (182 cells) and cells treated with iron for 2 hours (138 cells). Mean with errors bars representing SEM is indicated, n.s. indicates not significant as compared to control.

**Figure S3.**
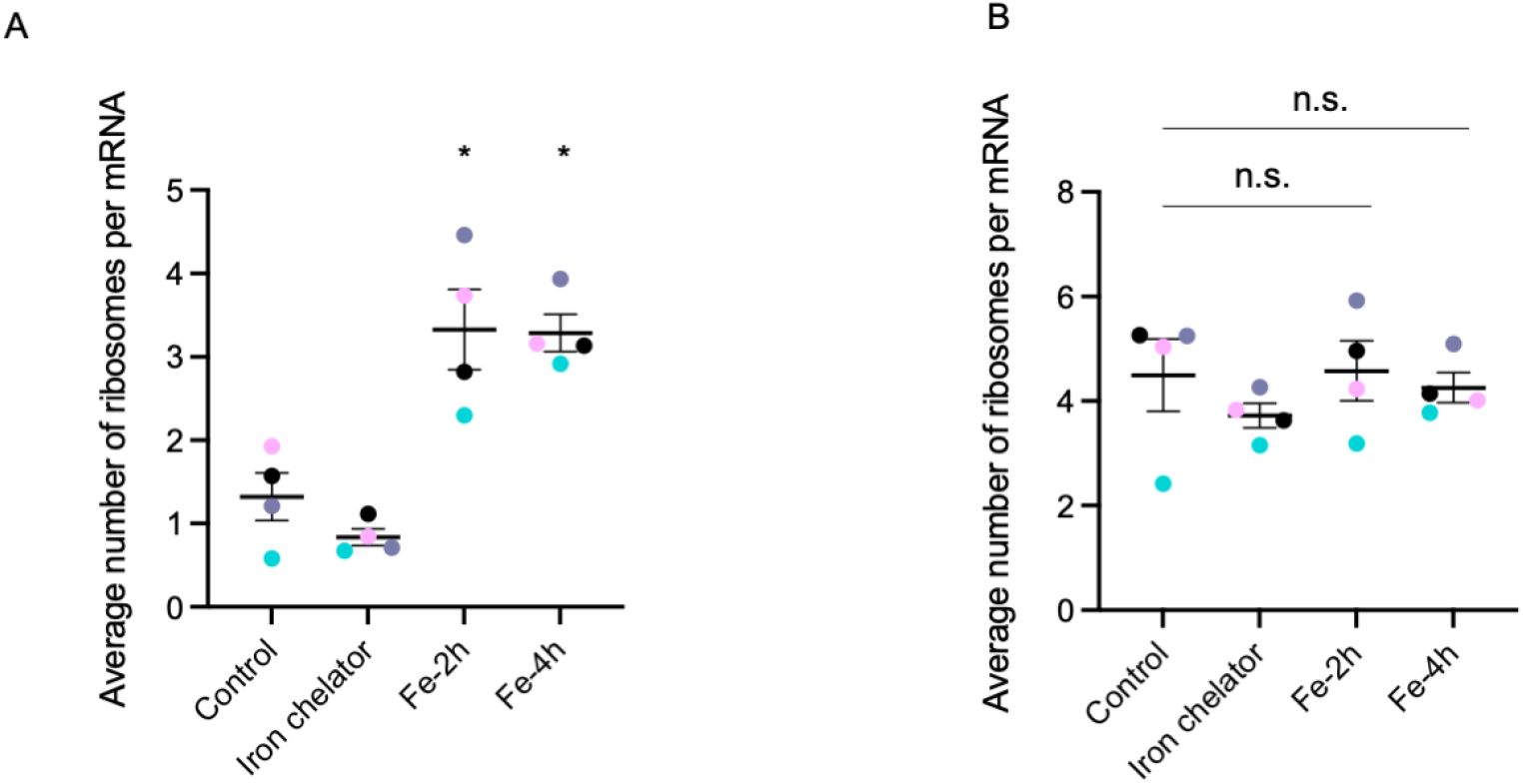
Quantification of ribosome occupancy on IRE-SunTag reporters. **A.** Average number of ribosomes per mRNA (for all mRNAs) in untreated control cells, iron chelator treated cells or cells treated with iron for 2h or 4h from 4 independent experiments. Color-coded dots represent the average number of ribosomes per mRNA in individual experiment sets. ^*^ Indicates p<0.05 as compared to control. The mean±SEM are represented. **B.** Average number of ribosomes per mRNA (for translating mRNAs only) in untreated control cells, iron chelator treated cells or cells treated with iron for 2h or 4h from 4-independent experiments. Color-coded dots represent the average number of ribosomes per mRNA in individual experiment sets. n.s. not significant when compared to control.

**Figure S4.**
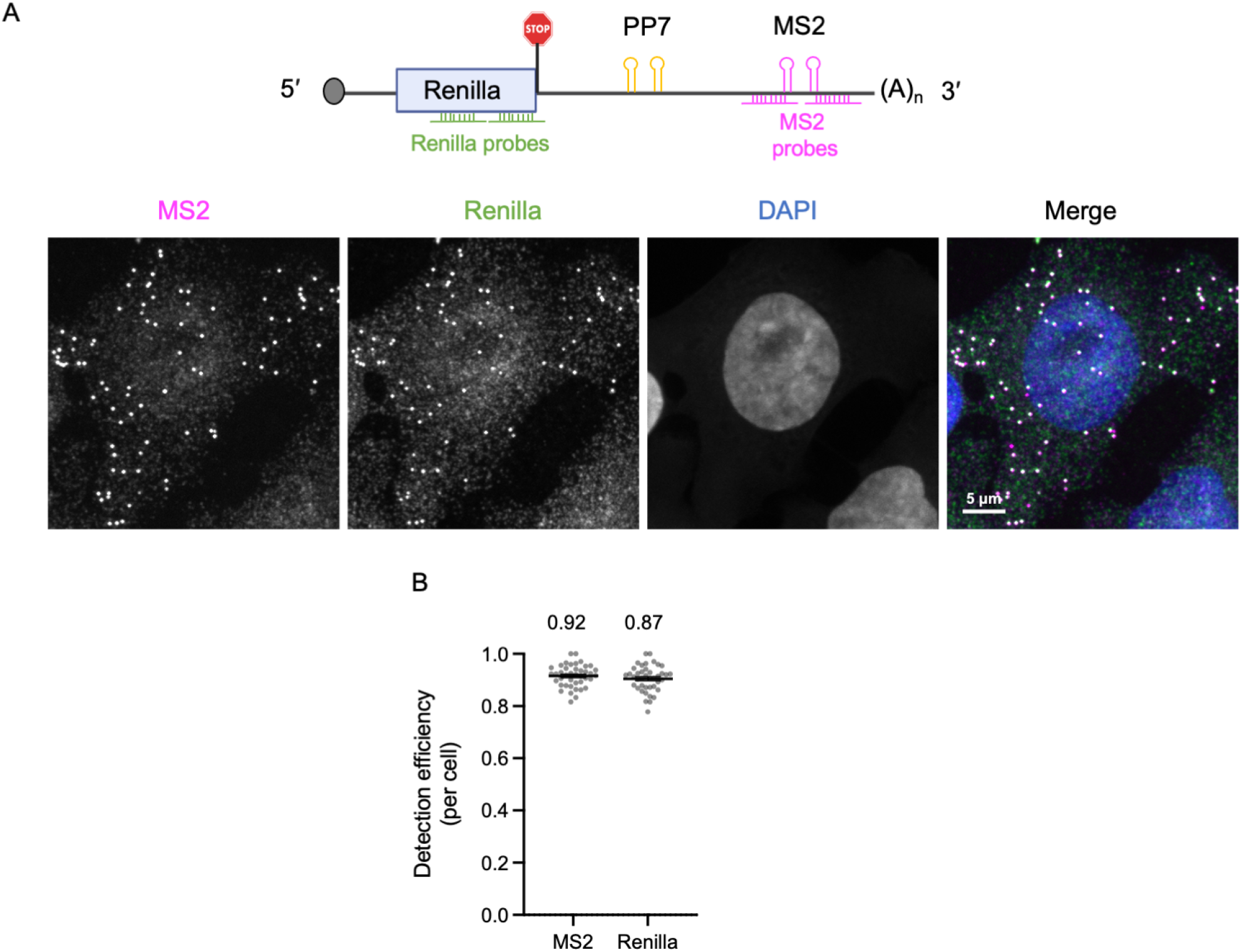
Determination of detection efficiencies of smFISH probes using minusPK control reporters. **A.** Schematic representation of the −pk-TREAT reporter, lacking the Xrn-1 resistant pseudoknots, used to quantify the detection efficiency of the probes. smFISH probes (Renilla and MS2) used for studying decay kinetics in all the experiments are represented. A representative smFISH image of a cell expressing −pk-TREAT reporter is presented. Magenta spots indicate signal from MS2 probes and green spots indicate signal from Renilla probes. **B.** The distribution of detection efficiency (see methods) of smFISH probes (MS2-Renilla probe pair) per cell. The mean detection efficiency MS2 probes and Renilla probes from all the cells is indicated above the plot.

**Figure S5.**
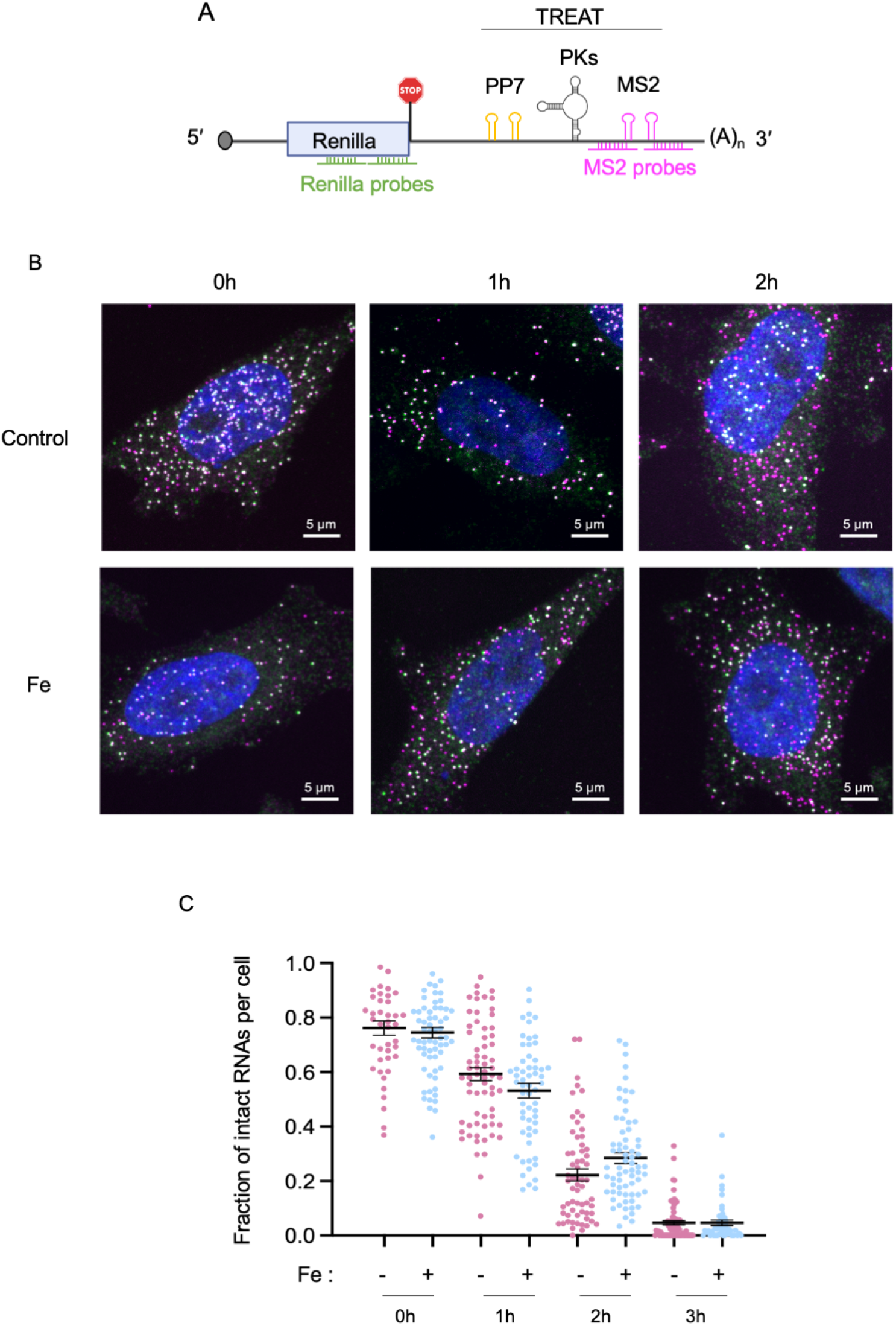
Iron-treatment does affect the decay of ΔIRE-TREAT reporter. **A.** Schematic representation of ΔIRE-TREAT reporter used to study translation upon treatment of cells with iron. smFISH probes targeting Renilla in ORF (upstream PKs) and MS2 stem-loops in 3’UTR (downstream PKs) is indicated. The reporter was integrated in HeLa cells under doxycycline inducible promoter. **B.** Representative smFISH images of cells expressing ΔIRE −TREAT reporters in presence or absence of iron at indicated time-points. Magenta spots represent MS2 and green spots represent Renilla. Dual labelled spots indicate the intact reporter RNAs and single labelled MS2 spots represent stabilized intermediates indicating degraded RNA. **C.** Quantification of mRNA decay from the smFISH images. Y-axis represents fraction of intact mRNAs corrected for the detection efficiency for the smFISH probes (see supplementary figure S4). Each dot represents fraction of intact mRNAs per cell. Mean and error bars indicating SEM is represented. Peach dots represent control and blue dots represent iron-treated cells.

**Figure S6.**
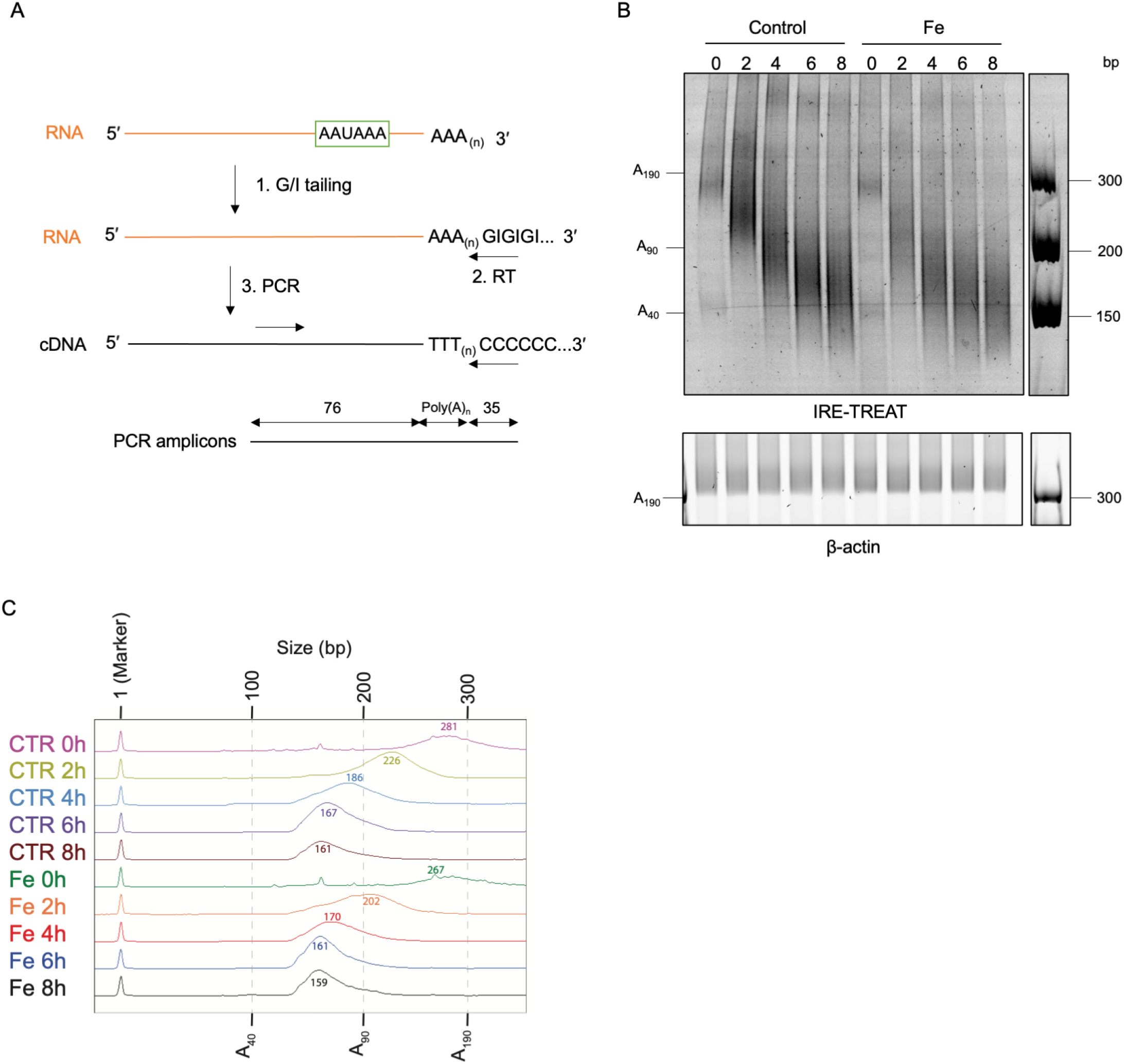
Poly(A) deadenylation of IRE-TREAT reporters. A. Schematic representation of the protocol used for qualitative assessment of the poly(A)n tail lengths of the IRE-TREAT reporter mRNA. Total RNA was isolated at time points indicated in figure 2B, and subsequently GI tails were added using poly(A)-polymerase followed by reverse transcription. Poly(A)n tails were amplified using a gene-specific forward primer and a universal reverse primer. B. The length of poly(A) tails in untreated and iron treated cells at indicated timepoints. Amplicon size as well as the calculated lengths of poly(A) tails is indicated. The poly(A)-tail lengths of steady state ß-actin mRNA was used as control. C. Quantification of the poly(A)-tail lengths using a Fragment Analyzer. PCR-products were purified and their size distribution was measured using a Fragment analyser. The curves reflect the distribution of the poly(A)-tail lengths, and size of the main peak is indicated above the curves in base pairs.

**Figure S7.**
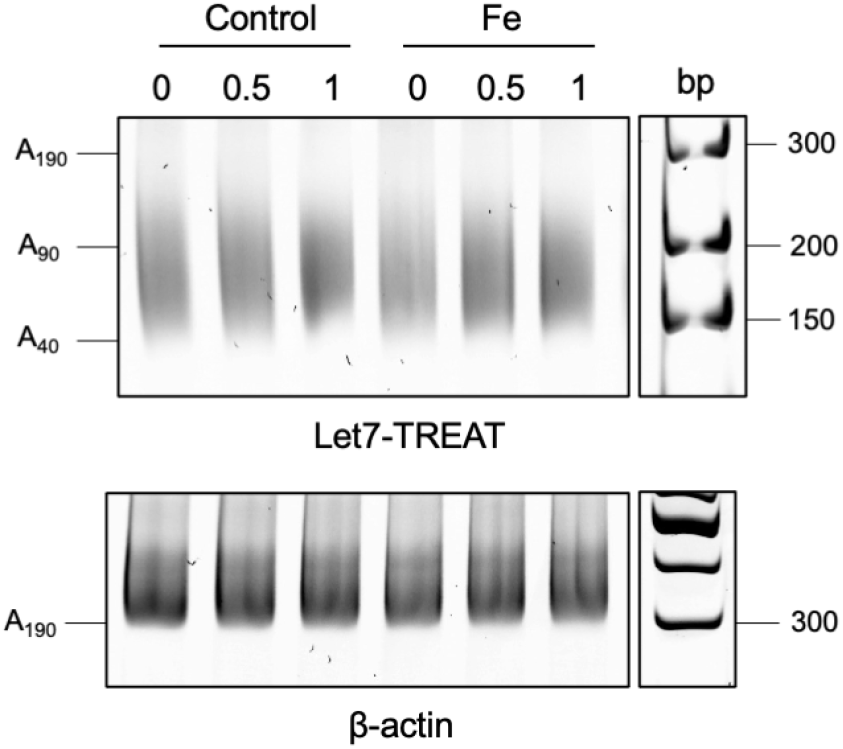
Deadenylation of miRNA regulated let7-TREAT reporters. The distribution of poly(A) tails of let7-TREAT reporter mRNA in untreated and iron-treated cells at indicated time points. Corresponding poly(A)-tail lengths are indicated.

**Figure S8.**
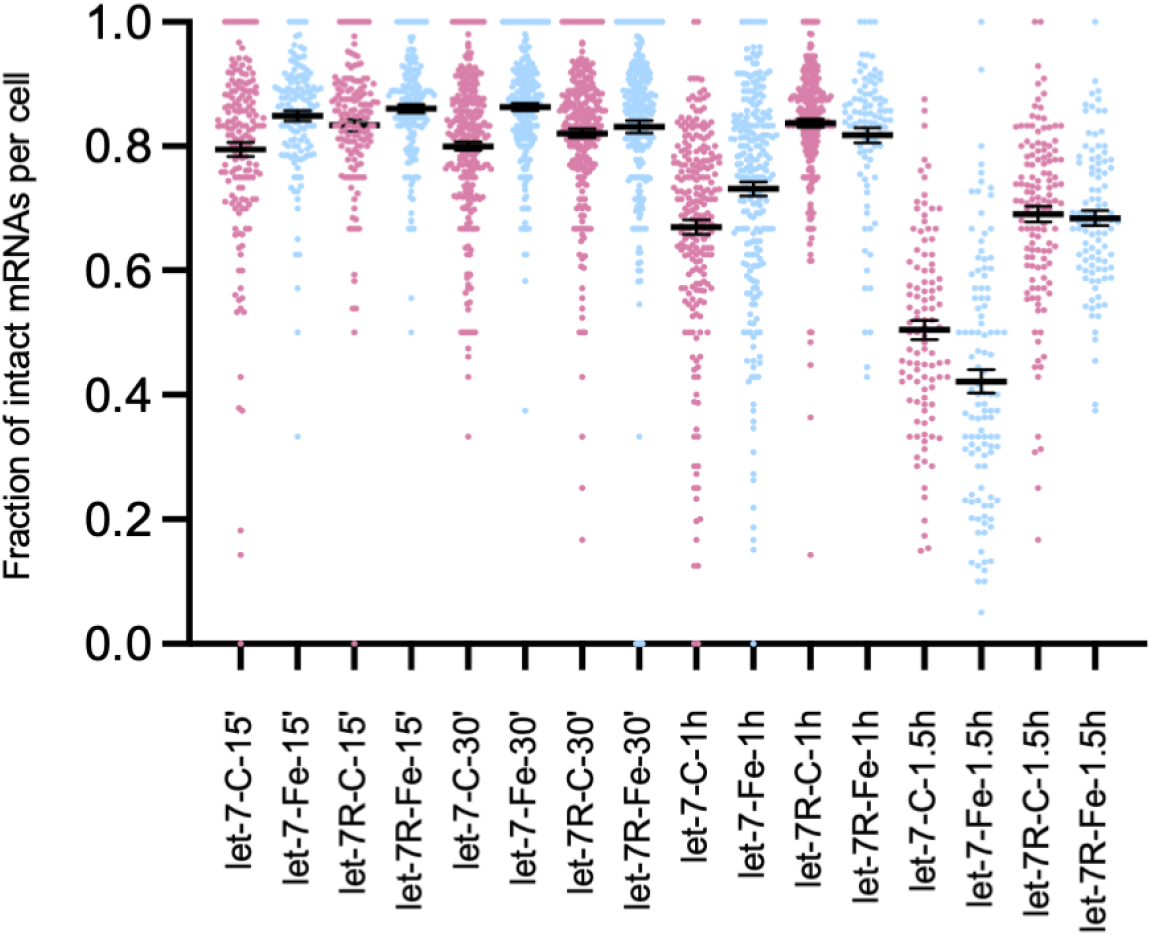
Quantification of fraction of intact mRNAs per cell for let7-SunTag-TREAT and let7R-SunTag-TREAT reporters, in untreated cells (peach spots) or iron-treated cells (blue spots). Each dot represents fraction of intact mRNAs per cell.

**Figure S9.**
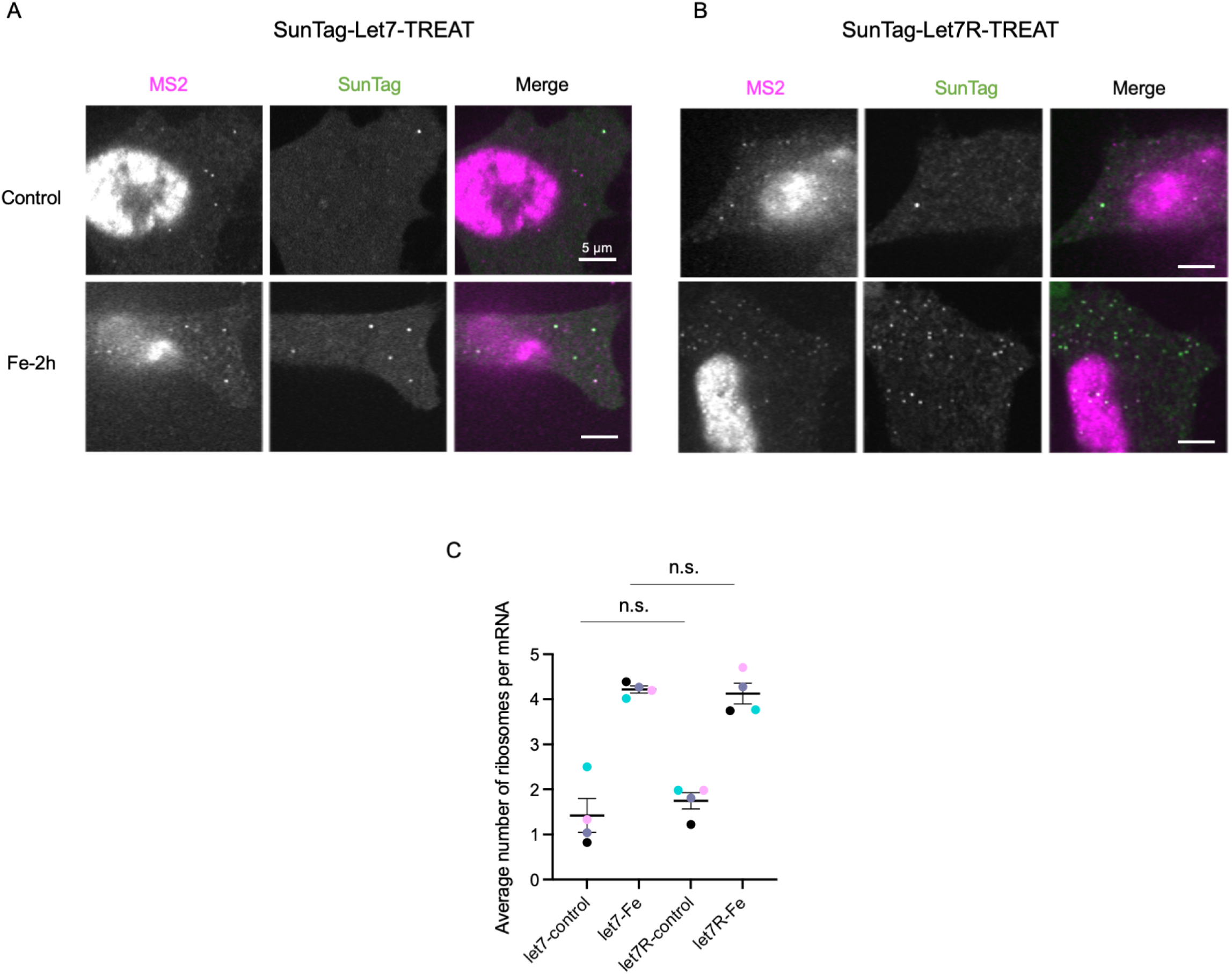
Live-cell imaging of let7-IRE-SunTag-TREAT and let7R-IRE-SunTag-TREAT reporters. **A. and B.** Live-cell single-molecule imaging of let7-SunTag-TREAT reporter (B) or let7R-SunTag-TREAT reporter (C) in untreated cells and iron-treated cells for 2 hours. The reporter mRNA was induced with doxycycline for 2 hours. Magenta spots represent mRNAs and green spots represents SunTag indicating translation. **B.** Average number of ribosomes per mRNA in untreated control cells, iron-treated cells expressing let7-SunTag-TREAT reporter or let7R-SunTag-TREAT reporter. Color-coded dots represent the average number of ribosomes per mRNA in individual experiment sets. n.s. not significant when compared to control.

## Supplementary video legends

**Video S1.** Translation of IRE-TREAT reporter mRNAs in absence of iron. Magenta spots indicate mRNAs (left) and green spots indicate SunTag (middle).

**Video S2.** Translation of IRE-TREAT reporter mRNAs in presence of iron treated for 2 hours. Magenta spots indicate mRNAs (left) and green spots indicate SunTag (middle).

**Video S3.** Translation of IRE-TREAT reporter mRNAs in presence of iron treated for 2 hours. Magenta spots indicate mRNAs (left) and green spots indicate SunTag (middle).

**Video S4.** Translation of IRE-TREAT reporter mRNAs upon treatment with 20 μM iron-chelator for 2 hours. Magenta spots indicate mRNAs (left) and green spots indicate SunTag (middle).

**Video S5.** Translation of ΔIRE-TREAT reporter mRNAs in absence of iron. Magenta spots indicate mRNAs (left) and green spots indicate SunTag (middle).

**Video S6.** Translation of ΔIRE-TREAT reporter mRNAs in presence of iron treated for 2 hours. Magenta spots indicate mRNAs (left) and green spots indicate SunTag (middle).

**Video S7.** Live cell imaging of mature SunTag peptides tethered to actin filaments in puromycin treated cells. Magenta spots indicate mRNAs (left) and green spots indicate SunTag peptides(middle). Arrows indicate stably associated mature SunTag peptides.

## Notes

### Competing Interest Statement

The authors have declared no competing interest.

## References

Banaszynski, L.A., Chen, L.C., Maynard-Smith, L.A., Ooi, A.G., and Wandless, T.J. (2006). A rapid, reversible, and tunable method to regulate protein function in living cells using synthetic small molecules. Cell 126, 995–1004.

Bartel, D.P. (2018). Metazoan MicroRNAs. Cell 173, 20–51.

Basham, B. (1997). Graphpad Prism. Biotechnol Softw I J 14, 14–17.

Beard, C., Hochedlinger, K., Plath, K., Wutz, A., and Jaenisch, R. (2006). Efficient method to generate single-copy transgenic mice by site-specific integration in embryonic stem cells. Genesis 44, 23–28.

Berman, A.J., Thoreen, C.C., Dedeic, Z., Chettle, J., Roux, P.P., and Blagden, S.P. (2021). Controversies around the function of LARP1. RNA Biol 18, 207–217.

Biasini, A., Abdulkarim, B., de Pretis, S., Tan, J.Y., Arora, R., Wischnewski, H., Dreos, R., Pelizzola, M., Ciaudo, C., and Marques, A.C. (2021). Translation is required for miRNA-dependent decay of endogenous transcripts. EMBO J 40, e104569.

Boersma, S., Khuperkar, D., Verhagen, B.M.P., Sonneveld, S., Grimm, J.B., Lavis, L.D., and Tanenbaum, M.E. (2019). Multi-Color Single-Molecule Imaging Uncovers Extensive Heterogeneity in mRNA Decoding. Cell 178, 458–472 e419.

Brengues, M., Teixeira, D., and Parker, R. (2005). Movement of eukaryotic mRNAs between polysomes and cytoplasmic processing bodies. Science 310, 486–489.

Buschauer, R., Matsuo, Y., Sugiyama, T., Chen, Y.H., Alhusaini, N., Sweet, T., Ikeuchi, K., Cheng, J., Matsuki, Y., Nobuta, R., et al. (2020). The Ccr4-Not complex monitors the translating ribosome for codon optimality. Science 368.

Chan, L.Y., Mugler, C.F., Heinrich, S., Vallotton, P., and Weis, K. (2018). Non-invasive measurement of mRNA decay reveals translation initiation as the major determinant of mRNA stability. Elife 7.

Chen, C.Y., Zheng, D., Xia, Z., and Shyu, A.B. (2009). Ago-TNRC6 triggers microRNA-mediated decay by promoting two deadenylation steps. Nat Struct Mol Biol 16, 1160–1166.

Cialek, C.A., Morisaki, T., Zhao, N., Montgomery, T.A., and Stasevich, T.J. (2021). Imaging translational control by Argonaute with single-molecule resolution in live cells. bioRxiv, 2021.2004.2030.442135.

Cottrell, K.A., Szczesny, P., and Djuranovic, S. (2017). Translation efficiency is a determinant of the magnitude of miRNA-mediated repression. Sci Rep-Uk 7.

Courel, M., Clement, Y., Bossevain, C., Foretek, D., Vidal Cruchez, O., Yi, Z., Benard, M., Benassy, M.N., Kress, M., Vindry, C., et al. (2019). GC content shapes mRNA storage and decay in human cells. Elife 8.

D’Orazio, K.N., Wu, C.C., Sinha, N., Loll-Krippleber, R., Brown, G.W., and Green, R. (2019). The endonuclease Cue2 cleaves mRNAs at stalled ribosomes during No Go Decay. Elife 8.

Dolken, L., Ruzsics, Z., Radle, B., Friedel, C.C., Zimmer, R., Mages, J., Hoffmann, R., Dickinson, P., Forster, T., Ghazal, P., et al. (2008). High-resolution gene expression profiling for simultaneous kinetic parameter analysis of RNA synthesis and decay. RNA 14, 1959–1972.

Eisen, T.J., Eichhorn, S.W., Subtelny, A.O., and Bartel, D.P. (2020a). MicroRNAs Cause Accelerated Decay of Short-Tailed Target mRNAs. Mol Cell 77, 775–785 e778.

Eisen, T.J., Eichhorn, S.W., Subtelny, A.O., Lin, K.S., McGeary, S.E., Gupta, S., and Bartel, D.P. (2020b). The Dynamics of Cytoplasmic mRNA Metabolism. Mol Cell 77, 786–799 e710.

Elkon, R., Zlotorynski, E., Zeller, K.I., and Agami, R. (2010). Major role for mRNA stability in shaping the kinetics of gene induction. BMC Genomics 11, 259.

Eraslan, B., Wang, D.X., Gusic, M., Prokisch, H., Hallstrom, B.M., Uhlen, M., Asplund, A., Ponten, F., Wieland, T., Hopf, T., et al. (2019). Quantification and discovery of sequence determinants of protein-per-mRNA amount in 29 human tissues. Mol Syst Biol 15.

Garneau, N.L., Wilusz, J., and Wilusz, C.J. (2007). The highways and byways of mRNA decay. Nat Rev Mol Cell Biol 8, 113–126.

Gaspar, I., Wippich, F., and Ephrussi, A. (2017). Enzymatic production of single-molecule FISH and RNA capture probes. RNA 23, 1582–1591.

Goossen, B., and Hentze, M.W. (1992). Position Is the Critical Determinant for Function of Iron-Responsive Elements as Translational Regulators. Molecular and Cellular Biology 12, 1959–1966.

Gowrishankar, G., Winzen, R., Dittrich-Breiholz, O., Redich, N., Kracht, M., and Holtmann, H. (2006). Inhibition of mRNA deadenylation and degradation by different types of cell stress. Biol Chem 387, 323–327.

Grimm, J.B., English, B.P., Chen, J., Slaughter, J.P., Zhang, Z., Revyakin, A., Patel, R., Macklin, J.J., Normanno, D., Singer, R.H., et al. (2015). A general method to improve fluorophores for live-cell and single-molecule microscopy. Nat Methods 12, 244–250, 243 p following 250.

Heck, A.M., and Wilusz, J. (2018). The Interplay between the RNA Decay and Translation Machinery in Eukaryotes. Cold Spring Harb Perspect Biol 10.

Hentze, M.W., Caughman, S.W., Rouault, T.A., Barriocanal, J.G., Dancis, A., Harford, J.B., and Klausner, R.D. (1987a). Identification of the iron-responsive element for the translational regulation of human ferritin mRNA. Science 238, 1570–1573.

Hentze, M.W., Rouault, T.A., Caughman, S.W., Dancis, A., Harford, J.B., and Klausner, R.D. (1987b). A cis-acting element is necessary and sufficient for translational regulation of human ferritin expression in response to iron. Proc Natl Acad Sci U S A 84, 6730–6734.

Herzog, V.A., Reichholf, B., Neumann, T., Rescheneder, P., Bhat, P., Burkard, T.R., Wlotzka, W., von Haeseler, A., Zuber, J., and Ameres, S.L. (2017). Thiol-linked alkylation of RNA to assess expression dynamics. Nature Methods 14, 1198-+.

Hilgers, V., Teixeira, D., and Parker, R. (2006). Translation-independent inhibition of mRNA deadenylation during stress in Saccharomyces cerevisiae. RNA 12, 1835–1845.

Hoek, T.A., Khuperkar, D., Lindeboom, R.G.H., Sonneveld, S., Verhagen, B.M.P., Boersma, S., Vermeulen, M., and Tanenbaum, M.E. (2019). Single-Molecule Imaging Uncovers Rules Governing Nonsense-Mediated mRNA Decay. Mol Cell 75, 324–339 e311.

Horvathova, I., Voigt, F., Kotrys, A.V., Zhan, Y., Artus-Revel, C.G., Eglinger, J., Stadler, M.B., Giorgetti, L., and Chao, J.A. (2017). The Dynamics of mRNA Turnover Revealed by Single-Molecule Imaging in Single Cells. Mol Cell 68, 615–625 e619.

Ibrahim, F., Maragkakis, M., Alexiou, P., and Mourelatos, Z. (2018). Ribothrypsis, a novel process of canonical mRNA decay, mediates ribosome-phased mRNA endonucleolysis. Nat Struct Mol Biol 25, 302–310.

Jackson, L.A., Anderson, E.J., Rouphael, N.G., Roberts, P.C., Makhene, M., Coler, R.N., McCullough, M.P., Chappell, J.D., Denison, M.R., Stevens, L.J., et al. (2020). An mRNA Vaccine against SARS-CoV-2 - Preliminary Report. N Engl J Med 383, 1920–1931.

Jia, L., Mao, Y., Ji, Q., Dersh, D., Yewdell, J.W., and Qian, S.B. (2020). Decoding mRNA translatability and stability from the 5’ UTR. Nat Struct Mol Biol 27, 814–821.

Juszkiewicz, S., Chandrasekaran, V., Lin, Z., Kraatz, S., Ramakrishnan, V., and Hegde, R.S. (2018). ZNF598 Is a Quality Control Sensor of Collided Ribosomes. Mol Cell 72, 469–481 e467.

Kawata, K., Wakida, H., Yamada, T., Taniue, K., Han, H., Seki, M., Suzuki, Y., and Akimitsu, N. (2020). Metabolic labeling of RNA using multiple ribonucleoside analogs enables the simultaneous evaluation of RNA synthesis and degradation rates. Genome Research 30, 1481–1491.

Kobayashi, H., and Singer, R.H. (2021). Single-molecule imaging of microRNA-mediated gene silencing in cells. bioRxiv, 2021.2004.2030.442050.

Kurosaki, T., Popp, M.W., and Maquat, L.E. (2019). Quality and quantity control of gene expression by nonsense-mediated mRNA decay. Nat Rev Mol Cell Biol 20, 406–420.

Laird-Offringa, I.A., de Wit, C.L., Elfferich, P., and van der Eb, A.J. (1990). Poly(A) tail shortening is the translation-dependent step in c-myc mRNA degradation. Molecular and Cellular Biology 10, 6132–6140.

Maroney, P.A., Yu, Y., Fisher, J., and Nilsen, T.W. (2006). Evidence that microRNAs are associated with translating messenger RNAs in human cells. Nature Structural & Molecular Biology 13, 1102–1107.

Mateju, D., Eichenberger, B., Voigt, F., Eglinger, J., Roth, G., and Chao, J.A. (2020). Single-Molecule Imaging Reveals Translation of mRNAs Localized to Stress Granules. Cell 183, 1801–1812 e1813.

Meijer, H.A., Kong, Y.W., Lu, W.T., Wilczynska, A., Spriggs, R.V., Robinson, S.W., Godfrey, J. D., Willis, A.E., and Bushell, M. (2013). Translational repression and eIF4A2 activity are critical for microRNA-mediated gene regulation. Science 340, 82–85.

Muckenthaler, M., Gray, N.K., and Hentze, M.W. (1998). IRP-1 binding to ferritin mRNA prevents the recruitment of the small ribosomal subunit by the cap-binding complex eIF4F. Mol Cell 2, 383–388.

Muckenthaler, M., Gunkel, N., Stripecke, R., and Hentze, M.W. (1997). Regulated poly(A) tail shortening in somatic cells mediated by cap-proximal translational repressor proteins and ribosome association. RNA 3, 983–995.

Otsu, N. (1979). Threshold Selection Method from Gray-Level Histograms. Ieee T Syst Man Cyb 9, 62–66.

Pelechano, V., Wei, W., and Steinmetz, L.M. (2015). Widespread Co-translational RNA Decay Reveals Ribosome Dynamics. Cell 161, 1400–1412.

Pichon, X., Bastide, A., Safieddine, A., Chouaib, R., Samacoits, A., Basyuk, E., Peter, M., Mueller, F., and Bertrand, E. (2016). Visualization of single endogenous polysomes reveals the dynamics of translation in live human cells. J Cell Biol 214, 769–781.

Pillai, R.S., Bhattacharyya, S.N., Artus, C.G., Zoller, T., Cougot, N., Basyuk, E., Bertrand, E., and Filipowicz, W. (2005). Inhibition of translational initiation by Let-7 MicroRNA in human cells. Science 309, 1573–1576.

Preibisch, S., Saalfeld, S., Schindelin, J., and Tomancak, P. (2010). Software for bead-based registration of selective plane illumination microscopy data. Nat Methods 7, 418–419.

Presnyak, V., Alhusaini, N., Chen, Y.H., Martin, S., Morris, N., Kline, N., Olson, S., Weinberg, D., Baker, K.E., Graveley, B.R., et al. (2015). Codon optimality is a major determinant of mRNA stability. Cell 160, 1111–1124.

Rueden, C.T., Schindelin, J., Hiner, M.C., DeZonia, B.E., Walter, A.E., Arena, E.T., and Eliceiri, K. W. (2017). ImageJ2: ImageJ for the next generation of scientific image data. BMC Bioinformatics 18, 529.

Santos, D.A., Shi, L., Tu, B.P., and Weissman, J.S. (2019). Cycloheximide can distort measurements of mRNA levels and translation efficiency. Nucleic Acids Res 47, 4974–4985.

Schindelin, J., Arganda-Carreras, I., Frise, E., Kaynig, V., Longair, M., Pietzsch, T., Preibisch, S., Rueden, C., Saalfeld, S., Schmid, B., et al. (2012). Fiji: an open-source platform for biological-image analysis. Nat Methods 9, 676–682.

Singh, M., Cornes, E., Li, B., Quarato, P., Bourdon, L., Dingli, F., Loew, D., Proccacia, S., and Cecere, G. (2020). Translation and codon usage regulate Argonaute slicer activity to trigger small RNA biogenesis. bioRxiv, 2020.2009.2004.282863.

Tat, T.T., Maroney, P.A., Chamnongpol, S., Coller, J., and Nilsen, T.W. (2016). Cotranslational microRNA mediated messenger RNA destabilization. Elife 5.

Tesina, P., Heckel, E., Cheng, J., Fromont-Racine, M., Buschauer, R., Kater, L., Beatrix, B., Berninghausen, O., Jacquier, A., Becker, T., et al. (2019). Structure of the 80S ribosome-Xrn1 nuclease complex. Nat Struct Mol Biol 26, 275–280.

Tinevez, J.Y., Perry, N., Schindelin, J., Hoopes, G.M., Reynolds, G.D., Laplantine, E., Bednarek, S.Y., Shorte, S.L., and Eliceiri, K.W. (2017). TrackMate: An open and extensible platform for single-particle tracking. Methods 115, 80–90.

Tuck, A.C., Rankova, A., Arpat, A.B., Liechti, L.A., Hess, D., Iesmantavicius, V., Castelo-Szekely, V., Gatfield, D., and Buhler, M. (2020). Mammalian RNA Decay Pathways Are Highly Specialized and Widely Linked to Translation. Mol Cell 77, 1222–1236 e1213.

Vogel, C., Abreu Rde, S., Ko, D., Le, S.Y., Shapiro, B.A., Burns, S.C., Sandhu, D., Boutz, D.R., Marcotte, E.M., and Penalva, L.O. (2010). Sequence signatures and mRNA concentration can explain two-thirds of protein abundance variation in a human cell line. Mol Syst Biol 6, 400.

Voigt, F., Gerbracht, J.V., Boehm, V., Horvathova, I., Eglinger, J., Chao, J.A., and Gehring, N.H. (2019). Detection and quantification of RNA decay intermediates using XRN1-resistant reporter transcripts. Nat Protoc 14, 1603–1633.

Voigt, F., Zhang, H., Cui, X.A., Triebold, D., Liu, A.X., Eglinger, J., Lee, E.S., Chao, J.A., and Palazzo, A.F. (2017). Single-Molecule Quantification of Translation-Dependent Association of mRNAs with the Endoplasmic Reticulum. Cell Rep 21, 3740–3753.

Wang, C., Han, B., Zhou, R., and Zhuang, X. (2016). Real-Time Imaging of Translation on Single mRNA Transcripts in Live Cells. Cell 165, 990–1001.

Weidenfeld, I., Gossen, M., Low, R., Kentner, D., Berger, S., Gorlich, D., Bartsch, D., Bujard, H., and Schonig, K. (2009). Inducible expression of coding and inhibitory RNAs from retargetable genomic loci. Nucleic Acids Res 37, e50.

Wilbertz, J.H., Voigt, F., Horvathova, I., Roth, G., Zhan, Y., and Chao, J.A. (2019). Single-Molecule Imaging of mRNA Localization and Regulation during the Integrated Stress Response. Mol Cell.

Wu, B., Eliscovich, C., Yoon, Y.J., and Singer, R.H. (2016). Translation dynamics of single mRNAs in live cells and neurons. Science 352, 1430–1435.

Wu, Q., Medina, S.G., Kushawah, G., DeVore, M.L., Castellano, L.A., Hand, J.M., Wright, M., and Bazzini, A.A. (2019). Translation affects mRNA stability in a codon-dependent manner in human cells. Elife 8.

Yamagishi, R., Hosoda, N., and Hoshino, S. (2014). Arsenite inhibits mRNA deadenylation through proteolytic degradation of Tob and Pan3. Biochem Biophys Res Commun 455, 323–331.

Yan, X., Hoek, T.A., Vale, R.D., and Tanenbaum, M.E. (2016). Dynamics of Translation of Single mRNA Molecules In Vivo. Cell 165, 976–989.

